# A DNA damage-induced phosphorylation circuit enhances Mec1^ATR^-Ddc2^ATRIP^ recruitment to Replication Protein A

**DOI:** 10.1101/2022.12.23.521831

**Authors:** Luke A. Yates, Elias A. Tannous, R. Marc Morgan, Peter M. Burgers, Xiaodong Zhang

## Abstract

The cell cycle checkpoint kinase Mec1^ATR^ and its integral partner Ddc2^ATRIP^ are vital for the DNA damage and replication stress response. Mec1-Ddc2 ‘senses’ single-stranded DNA (ssDNA) by being recruited to the ssDNA binding Replication Protein A (RPA) via Ddc2. In this study, we show that a DNA-damage induced phosphorylation circuit modulates checkpoint recruitment and function. We demonstrate that Ddc2-RPA interactions modulate the association between RPA and ssDNA and that Rfa1-phosphorylation aids in the further recruitment of Mec1-Ddc2. We also uncover an underappreciated role for Ddc2 phosphorylation that enhances its recruitment to RPA-ssDNA that is important for the DNA damage checkpoint in yeast. The crystal structure of a phosphorylated Ddc2 peptide in complex with its RPA interaction domain provides molecular details of how checkpoint recruitment is enhanced, which involves Zn^2+^. Using electron microscopy and structural modelling approaches, we propose that Mec1-Ddc2 complexes can form higher order assemblies with RPA when Ddc2 is phosphorylated. Together, our results provide insight into Mec1 recruitment and suggest that formation of supramolecular complexes of RPA and Mec1-Ddc2, modulated by phosphorylation, would allow for rapid clustering of damage foci to promote checkpoint signalling.

**Graphical Abstract:** 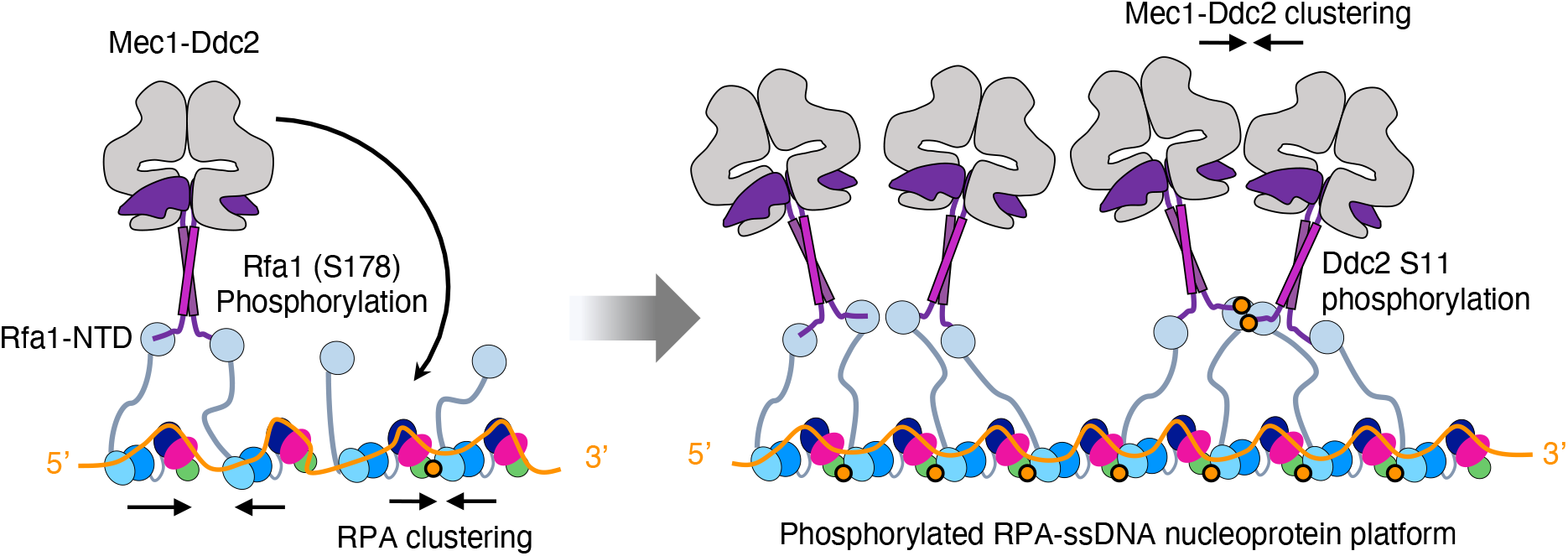

**Highlights:**

- Rfa1-S178 phosphorylation promotes Ddc2 recruitment and Ddc2-RPA complexes modulate RPA-ssDNA behaviour.
- Ddc2 phosphorylation enhances Mec1-Ddc2 recruitment and is important for the DNA damage checkpoint in yeast.
- Structure of a Ddc2:RPA complex shows phosphorylation-dependent higher order assemblies stabilised by Zn^2+^.
- We propose a Mec1-Ddc2 recruitment strategy that allows fast accumulation of Mec1-Ddc2 through DNA damage-induced phosphorylation and promotes autophosphorylation.

## Introduction

The cell cycle checkpoint is a signalling pathway that coordinates DNA repair and replication challenges with cell cycle progression to safeguard the genome. The kinase Mec1 (Mitosis Entry Checkpoint), and its human counterpart ATR (ATM and Rad3 related), are master regulators of the checkpoint that are crucial for DNA replication, DNA damage, and replication stress responses. Mec1, a member of the Phosphatidylinositol 3-kinase like kinase (PIKK) family, is essential in yeast (Kato and Ogawa, 1994; Weinert et al., 1994) and ATR loss is early embryonic lethal in mammals (Brown and Baltimore, 2000; Liu et al., 2000). Mec1/ATR dysfunction increases genomic instability in S-phase and cause chromosomal rearrangements (Buisson et al., 2015; Casper et al., 2002). Hypomorphic mutations in human ATR is associated with Seckel syndrome, an autosomal recessive disorder characterised by microencephaly, dwarfism, and intellectual disability (O’Driscoll et al., 2003).

Double strand breaks occurring in S and G2 phase are repaired by homologous recombination (HR), which utilises regions of homology in sister chromatids as a template. During HR, the broken DNA ends are resected to produce long 3’ single-stranded DNA (ssDNA) overhangs. Stalled replication forks also produce long and persistent stretches of ssDNA. In both cases the ssDNA is coated by RPA, a ubiquitous ssDNA binding protein that protects the DNA from degradation, undesired pairing, and secondary structure formation (Wold, 1997). Mec1/ATR is recruited to RPA-ssDNA nucleoprotein filaments via its integral binding partner Ddc2/ATRIP (Zou and Elledge, 2003). Mec1/ATR coordinates cell cycle progression with DNA repair or fork stabilisation and a major cellular consequence of its activity is histone H2A phosphorylation (a bona fide signal of DNA damage) and cell cycle arrest via Rad53/CHK1 phosphorylation. Mec1-Ddc2 therefore maintains a low level of kinase activity, which is not increased upon RPA-dependent recruitment, but instead requires cell-cycle specific activators for proper function (Tannous and Burgers, 2021). We recently uncovered a mechanism for how Mec1-Ddc2 is maintained in an auto-inhibited state, and the conformational changes required to transition into an activated state (Tannous et al., 2021). Given that a number of Mec1 activators are associated with chromatin, controlled recruitment of Mec1-Ddc2 by post-translational modifications (PTMs) could also be a means to regulate its cellular activity (Dubois et al., 2017; Maréchal et al., 2014; Memisoglu et al., 2019). Autophosphorylation of Mec1 is also important for its function (Hurst et al., 2021; Sanford et al., 2021) and occurs *in trans* (Memisoglu et al., 2019) implying that Mec1-Ddc2 must be brought together for this to occur. The localised Mec1 response at the DNA lesion site, characterised by H2A S129 phosphorylation, is rapid and hypersensitive to low levels of ssDNA (Bantele et al., 2019). Further, Mec1-Ddc2 is anchored at specific locations in damage chromatin (Bantele et al., 2019), suggesting a DNA-damage induced mechanism for rapid and stable recruitment of Mec1-Ddc2 to RPA-ssDNA, which we hypothesise relies on post-translation modifications (PTMs). However, a detailed molecular understanding of how Mec1-Ddc2 recruitment to RPA-ssDNA could be modulated by PTMs remains unclear.

Yeast RPA consists of three subunits, Rfa1 (RPA70 in human), Rfa2 (RPA32) and Rfa3 (RPA14), with each subunit consisting of one or more ssDNA-binding OB-fold domains (DBDs). The Rfa1/RPA70 N-terminal domain (Rfa1-NTD) is a predominantly protein-interacting module and specifically interacts with Ddc2/ATRIP (Haring et al., 2008). Previous studies have shown that direct interactions between Rfa1-NTD and the N-terminal region of Ddc2 (Ball et al., 2007) or ATRIP (Choi et al., 2010) are essential for checkpoint function. Structural studies of the Mec1-Ddc2, which is a stable dimer of heterodimers (Tannous et al., 2021), together with a complex structure between a homodimeric N-terminal coiled-coil domain of *K. lactis* Ddc2 and *Saccharomyces cerevisiae* Rfa1-NTD (Deshpande et al., 2017), suggests a 2:2:2 arrangement of RPA, Mec1 and Ddc2. Further, the N-terminal Ddc2 α-helical acidic domain is engulfed by a positively charged groove of the Rfa1-NTD OB-fold, similar to other Rfa1/RPA70-NTD interacting domains (Deshpande et al., 2017; Oakley and Patrick, 2010). Additionally, RPA interacts with a many repair factors through the Rfa1/RPA70-NTD via its positively charged groove, raising questions about how different RPA-interacting DNA damage factors are selectively recruited. Previously, we have shown that *Saccharomyces cerevisiae* RPA forms oligomers on ssDNA via an inter-RPA interaction between the OB domain of Rfa1 and the OB domain of Rfa3 of an adjacent RPA. Importantly, we have shown that this oligomerisation is enhanced using a phosphomimetic substitution at S178 of the Rfa1 subunit (Yates et al., 2018), which is a Mec1 dependent phosphorylation site (Bastos de Oliveira et al., 2015). However, the precise effect of this phosphorylation on the recruitment of Mec1-Ddc2 and damage foci formation remain unclear.

In this work, we set out to investigate the interactions between RPA and Mec1-Ddc2 and the effect of site-specific phosphorylation on kinase recruitment. Our results reveal that Rfa1-S178 phosphorylation enhances Mec1-Ddc2 recruitment, and the binding of Ddc2 dimers in turn enhances RPA clustering. Further, we find that Serine11 phosphorylation in Ddc2 modulates the interactions between Mec1-Ddc2 and RPA and plays a crucial role in Mec1-Ddc2 dependent DNA damage response. Significantly, our studies uncover a novel mode of protein oligomerisation through phosphorylation, with a role for Zn^2+^ binding within this assembly. We propose a recruitment strategy that allows fast accumulation of Mec1-Ddc2 at damaged sites through DNA damage-induced phosphorylation of RPA and Ddc2, potentially stabilised by Zn^2+^ ions, that has the potential to promote damage foci and Mec1 autophosphorylation.

## Results

### Phosphorylation of RPA at S178 enhances Mec1-Ddc2 recruitment

We previously provided a model for how multiple RPAs form higher order structures to coat ssDNA (Yates et al., 2018). We also showed that mimicking phosphorylation of Rfa1-S178 causes enhanced *trans* RPA-RPA interactions resulting in cooperative ssDNA binding (Yates et al., 2018). RPA is a major platform for Mec1-Ddc2 recruitment (Dubrana et al., 2007; Rouse and Jackson, 2002; Zou and Elledge, 2003), and the removal of RPA from ssDNA via the Srs2 helicase, for example, has been shown to dampen the checkpoint (Dhingra et al., 2021). Therefore, we reasoned that the enhanced clustering of RPA via Rfa1 phosphorylation could play a role in checkpoint kinase recruitment. We set out to address this question by characterising interactions between Mec1-Ddc2 with RPA alone and bound to ssDNA. Previous work has established that the Mec1-Ddc2 complex forms head-to-head dimers of Mec1-Ddc2 heterodimers via several regions, including the Mec1 kinase, FAT domains, and Ddc2 C-terminal HEAT repeats (**Figure 1A**) (Tannous et al., 2021; Wang et al., 2017). Further, the N-terminal region of Ddc2 forms a coiled-coil dimer and, importantly, possesses a well-characterised RPA-interaction site (**Figure 1A**) (Ball et al., 2007; Deshpande et al., 2017). Using biotinylated-dT_32_ oligonucleotide immobilised to streptavidin beads we were able to show that phosphomimetic RPA^S178D^ (RPA with an Rfa1^S178D^ subunit) binding to purified Mec1-Ddc2 complexes was not grossly perturbed compared to RPA^WT^ (**Supplementary Figure 1**). Due to the limited quantity of purified Mec1-Ddc2, and given that the major RPA-interaction sites are situated at the N-terminus of Ddc2, we constructed and purified an N-terminal Ddc2 fragment containing the main RPA binding site together with the coiled-coil domain (residues 1-148, denoted as Ddc2-CC, **Figure 1A-B**), which should maintain a dimeric state. Indeed Ddc2-CC domains are dimers, confirmed using SEC-MALS, and are able to interact with RPA, measured by microscale thermophoresis (MST) (**Supplementary Figure 2A-B**). Using this Ddc2-CC fragment, we assessed its binding to RPA coated ssDNA using MST. Ddc2-CC binds to RPA-dT100 nucleoprotein with similar affinity to RPA alone (**Figure 1B and Supplementary Figure 2C**). RPA^S178D^ showed an improved affinity (2-fold) for the Ddc2-CC fragment (**Figure 1B**), suggesting that Rfa1^S178^ phosphorylation enhances Ddc2 interactions, and by extension Mec1-Ddc2 recruitment, which supports yeast co-immunoprecipitation experiments with RPA-S178A that show a 50% reduced Mec1 recruitment capacity (Kim and Brill, 2003).

**Figure 1.**
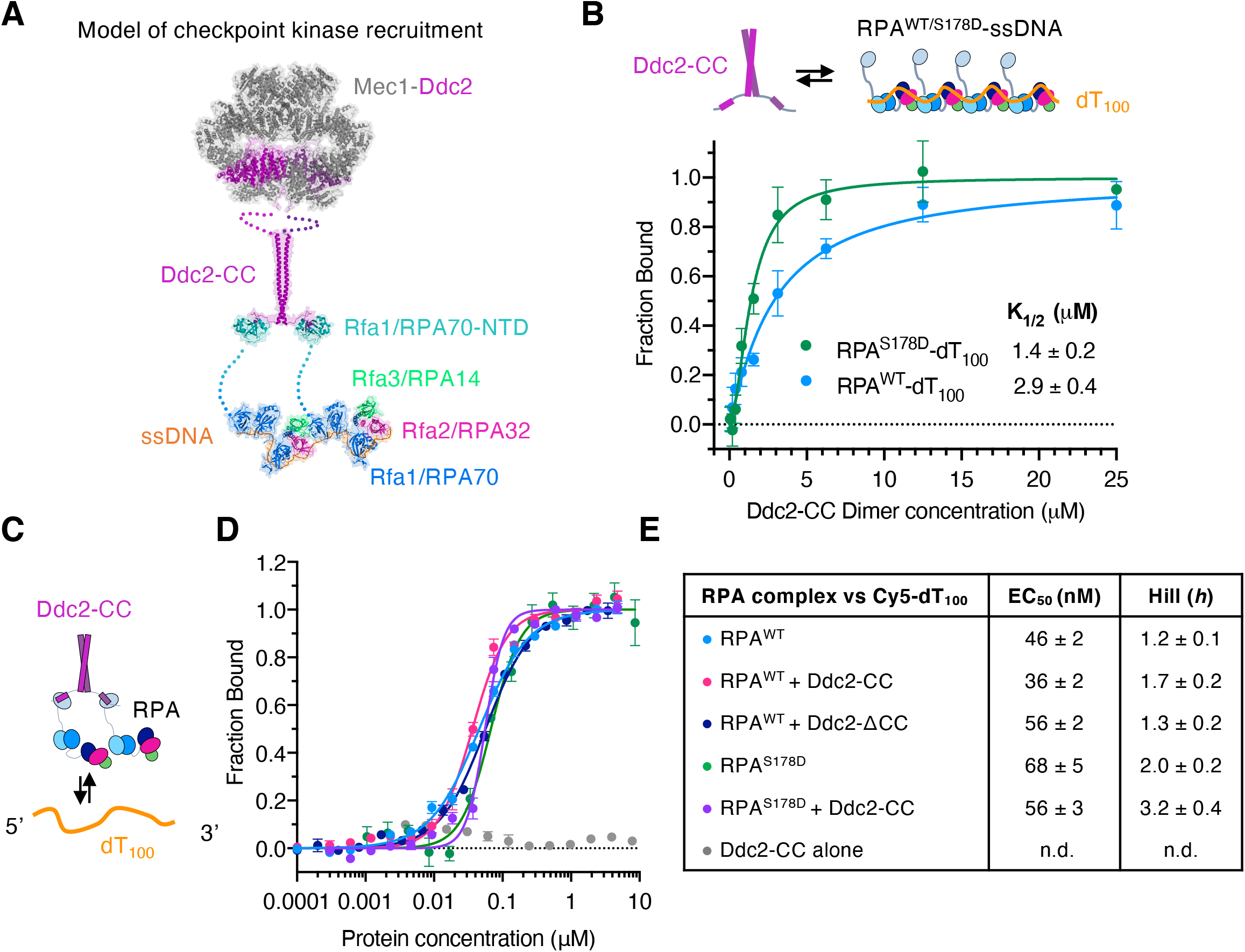
Interaction studies between Mec1-Ddc2 and RPA. **A** Model of Mec1-Ddc2 recruited by RPA-ssDNA based upon available structural data (Deshpande et al., 2017; Tannous et al., 2021; Yates et al., 2018). **B** Microscale thermophoresis (MST) binding curves of Ddc2-CC versus RPA_^WT^_ or phosphomimetic mutant RPA (RPA^S178D^) bound to homopolymer ssDNA (dT_100_). **C** Schematic of Ddc2-CC dimers tethering RPA to influence its association with ssDNA. **D-E** MST-binding curves of RPA complexes interacting with Cy5-labeled dT_100_ ssDNA substrate, using RPA^WT^ or RPA^S178D^, in the absence or presence of Ddc2-CC. For all MST curves, apparent EC_50_ and Hill values were calculated using the Hill-equation and tabulated (e). n.d. indicates not determined. Normalized fluorescence were calculated using NanoTemper Analysis 1.2.101. Measurements were repeated at least 3 times or more and data points are averages with standard error (SE).

We next tested whether Ddc2-CC:RPA complexes alter the behaviour of RPA nucleoprotein filament assembly. To quantify these interactions, we used MST to measure direct binding between RPA and ssDNA in the absence and presence of Ddc2-CC. The addition of Ddc2-CC does not affect the interaction between RPA and ssDNA (**Figure 1E**). This is not surprising, as Ddc2-CC does not bind to ssDNA (**Figure 1E**) but interact with Rfa1-NTD, which is independent of the DNA binding domains (Brosey et al., 2015). However, the tethering of RPAs via the homodimeric Ddc2 fragment causes ssDNA binding by RPA to become co-operative, as suggested by the Hill coefficient, but not when the coiled-coil region is removed (**Figure 1E**). This cooperativity is further enhanced when combined with the RPA^S178D^ mutant, with hill values suggesting an additive effect due to two independent mechanisms coupling RPA molecules together. Additional experiments using electrophoretic mobility shift assays (EMSA) show improved RPA coating in the presence of Ddc2-CC, consistent with the cooperativity observed by MST (**Supplementary Fig 2D**). Further, Ddc2-CC domain can also bridge two separate RPA coated ssDNA molecules together (**Supplementary Figure 2E-F**), which implies that Mec1-Ddc2 can be recruited to separate regions of RPA-ssDNA that are close in space, such as RPA-ssDNA loops caused by replication (Burgers, 2009), or during end-resection (Xue et al., 2022), for example.

Collectively, these data show that *trans* RPA-RPA interactions that promote clustering are important for the recruitment of Mec1-Ddc2 complexes. In addition, the tethering of RPA via its interaction with dimeric Ddc2 promotes cooperative binding to ssDNA, with a possible role in bridging distal regions of RPA-ssDNA or at RPA-ssDNA loops. This synergistic enhancement results in more stably recruited Mec1-Ddc2 complexes and infers a feed-forward loop involving phosphorylation of Rfa1, which is predominantly Mec1-dependent (Bastos de Oliveira et al., 2015), in recruiting and retaining checkpoint kinase at ssDNA.

### Phosphorylation of the Ddc2 N-terminus enhances RPA association

Since Rfa1-S178 phosphorylation influences Mec1-Ddc2 association, we next asked if phosphorylation of Mec1-Ddc2 can also influences recruitment. Proteomic and genetic studies have demonstrated a number of phosphorylation sites in Mec1-Ddc2 that are important for function (Hustedt et al., 2015; Memisoglu et al., 2019). We found that purified Mec1-Ddc2 used in our previous studies is extensively phosphorylated and the majority of Ddc2’s phosphorylation can be removed by phosphatase treatment (**Supplementary Figure 3A**). The Mec1 phosphosites previously described (Memisoglu et al., 2019) are not easily accessible to phosphatase treatment, and we suspect that these are buried in this functional state. This prompted us to investigate the role of Mec1-Ddc2 phosphorylation on RPA-ssDNA interaction. We purified Mec1-Ddc2 complexes in which Ddc2 harboured a FLAG-tag at the N-terminus (a kind gift of Prof. J. Diffley) (**Supplementary Figure 3B**) and labelled it with a fluorescent (Alexa Fluor-647) anti-FLAG antibody. During the Mec1-Ddc2 purification, we carefully preserved the phosphorylation status by including phosphatase inhibitors at every step. Finally, we separated the Mec1-Ddc2 samples into two aliquots; one treated with lambda-phosphatase, and the other with reaction buffers only, resulting in phosphorylated or dephosphorylated samples (**Figure 2A**). MST binding results using a titration of RPA-dT_100_ nucleoprotein complexes produced a dissociation constant of 0.7 μM for phosphorylated Mec1-Ddc2 (**Figure 2B**). Dephosphorylated Mec1-Ddc2 produced a Kd of 4.1 μM (**Figure 2B**), ∼6-fold weaker than phosphorylated samples but is similar to the Kd obtained for unmodified Ddc2-CC (**Supplementary Figure 2C**). These data suggest that phosphorylation of Mec1-Ddc2 (most likely Ddc2) enhances binding to RPA-ssDNA.

**Figure 2.**
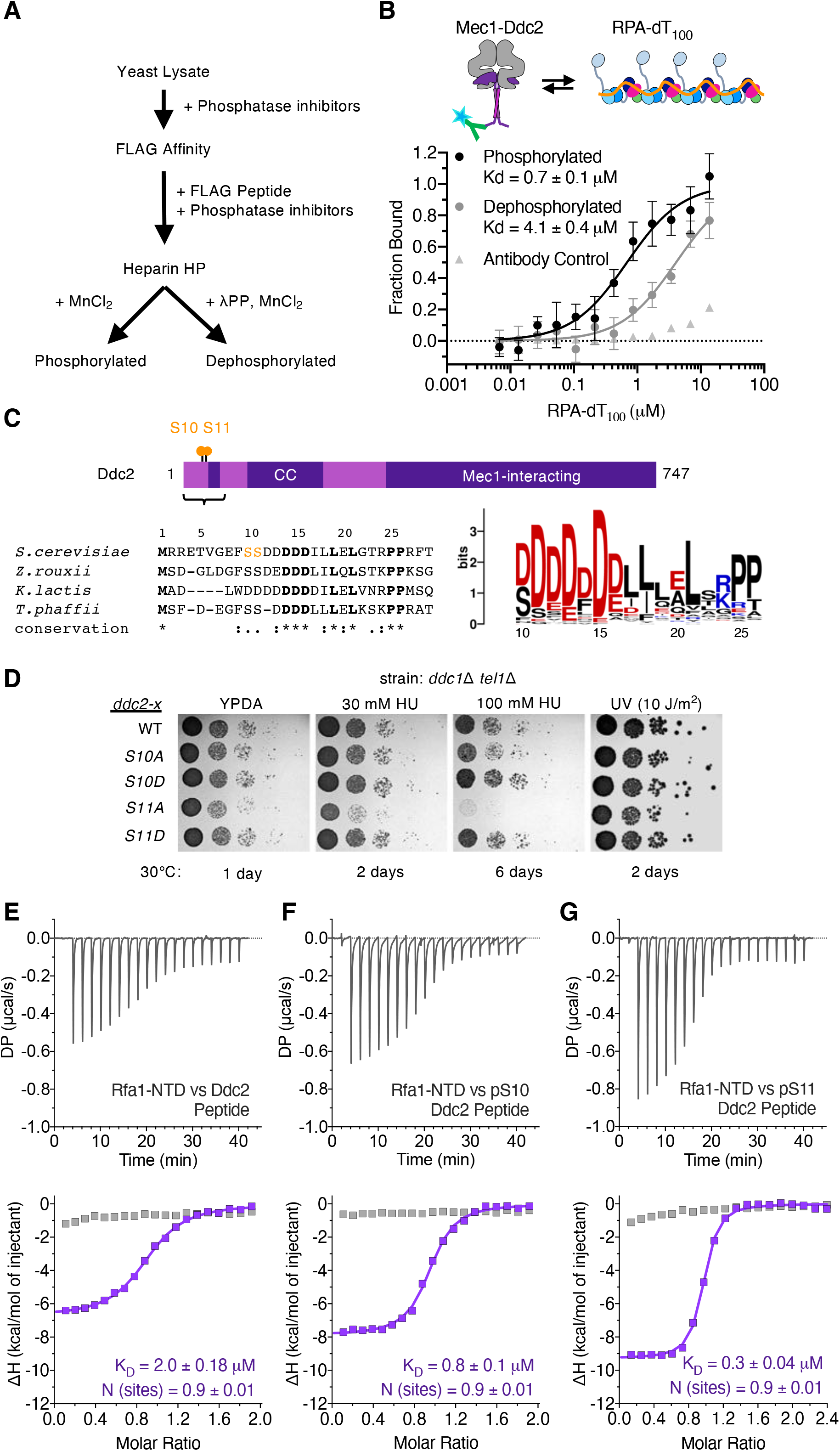
Phosphorylation of Ddc2 enhances recruitment to RPA and is important for the damage response. **A** Purification strategy for comparing phosphorylated and dephosphorylated (treated with λ-protein phosphatase, λPP) Mec1-Ddc2 by MST. **B** MST binding curves using Mec1-Ddc2 or Mec1-Ddc2 treated with lambda-phosphatase vs RPA-ssDNA. Approximately 100nM Mec1-Ddc2 complex was mixed with equimolar amounts of Alexafluor-647 conjugated anti-FLAG antibody to label the Ddc2 subunit, which harbours an N-terminal 3xFLAG tag, and used for MST versus a serial dilution of RPA-ssDNA at concentrations indicated. Antibody alone was used as a control. Fraction bound was estimated using the averaged normalised fluorescence and fitting the data using a law of mass action equation in Nanotemper analysis software. **C** schematic of domains in Ddc2 along with its phosphorylation sites. S10 and S11, the focus of this study, are shown by orange circles. A sequence alignment of the RPA-interacting region from other yeast species is shown underneath. An amino acid frequency plot from alignment of 90 Ddc2 sequences from different fungi species showing a high conservation of acidic residues. Numbering underneath corresponds to *S. cerevisiae* Ddc2 numbering. **D** Growth of *DDC2* mutants. Strain PY437 is *ddc2Δ tel1Δ ddc1Δ* and contain plasmid p(*LEU2 ddc2-x*). The strain was tested for growth on YPDA plates with or without hydroxyurea or treated with UV as indicated. **E-G** Raw Isothermal Titration Calorimetry (ITC) (upper panels) and normalised heat of binding isotherms (lower panels) for Ddc2 peptides and phosphorylation variants titrated against Rfa1-NTD (purple) or buffer alone (heat of dilution controls, grey). Calculated dissociation constants (K_D_) are derived from the measured association constant by linear regression fitting using a 1-to-1 binding model in MicroCal PEAQ-ITC analysis software.

To identify potential phosphorylation sites responsible for the enhanced binding to RPA, we focused on those phosphosites previously described in Ddc2 that are close to the known RPA interaction sites – a motif that is often identified by a negatively charged Asp-rich region (Lee et al., 2020). Proteomic studies have identified two Ddc2 phosphorylation sites, Serine10 and Serine11, adjacent to the highly conserved RPA-interacting motif (Albuquerque et al., 2008; Holt et al., 2009; Lanz et al., 2021) (**Figure 2C**). Both Serine10 and Serine11 phosphorylation have been robustly detected when cells are exposed to methylmethane sulfonate (MMS) (Zhou et al., 2016). These two phosphosites are also conserved in other species but are often found to be substituted by Aspartate, particularly at the equivalent S11 site, suggesting natural phosphomimetics (**Figure 2C**). We thus introduced substitutions into Ddc2 to mimic states that are constitutively phosphorylated (Ser to Asp) or unphosphorylated (Ser to Ala) and tested there effect on growth and sensitivity to replication inhibitor hydroxyurea (HU) or UV damage in several indicator strains that are progressively compromised for the cell cycle checkpoint circuitry (**Figure 2D** and **Supplementary Figure 4A**). Ddc2-S10A, Ddc2-S10D, and Ddc2-S11D phenocopies WT yeast under the conditions tested (**Figure 2D**). On the other hand, S11A shows growth defects and is very sensitive to HU, but limited UV sensitivity (**Figure 2D**). This sensitivity was also phenocopied by a S10A-S11A double mutant, whereas S10D-S11D mutants are analogous to WT (**Supplementary Figure 4A**), suggesting non-phosphorylatable S11 compromises Ddc2 function in replication stress. Kinase activity of purified Mec1-Ddc2(S10D/A) and Mec1-Ddc2(S11D/A) mutants were not obviously compromised, suggesting the checkpoint defective phenotypes observed are related to recruitment deficiencies in yeast (**Supplementary Figure 4B**). We therefore predicted that both S10 and S11 phosphosites in Ddc2 could modulate the association with RPA, given their close proximity to the known interaction motif. To this end, we tested the relative affinities between synthetic phosphorylated and unmodified Ddc2 peptides (amino acids 4-24) with Rfa1-NTD (residues 1-132) using isothermal titration calorimetry (ITC) (**Figure 2E-G, Supplementary Figure 4C**). ITC isotherms show that the unmodified Ddc2 peptide (denoted WT) binds to Rfa1-NTD with an ∼2 μM dissociation constant (KD) (**Figure 2E**). Phosphorylation of either Serine10 (pS10) or Serine11 (pS11) produced approximately 3-fold and 7-fold improved affinity, respectively (**Figure 2F-G**). A bis-phosphorylated peptide (pS10-pS11) showed a similar affinity to pS11-peptide, suggesting S11 plays a dominant role (**Supplementary Figure 4C**). Interestingly, the fold increase in affinity observed for S11 phosphorylation is similar to that observed between phosphorylated and dephosphorylated Mec1-Ddc2 binding to RPA-ssDNA. This is consistent with the idea that phosphorylation of Ddc2 could be responsible for the enhanced binding between Mec1-Ddc2 and RPA-ssDNA. Our genetic studies along with our interaction studies show that S11 is more important for function *in vivo*, consistent with ITC data that S11 phosphorylation has a larger effect on binding (**Figure 2F-G**).

### Structure of Rfa1-NTD in complex with a phosphorylated Ddc2 peptide

To understand the molecular basis of the phosphorylation events and how it might affect the binding, we determined a crystal structure of *Sc*Rfa1-NTD in complex with a phosphorylated *Sc*Ddc2 peptide (residues 4-24) to 1.58 Å resolution (**Figure 3A-D, Table 1, and Supplementary Figure 5A-C**). The structure was solved by molecular replacement using the Rfa1-NTD structure from (PDB:5OMB) as a search model. We were able to clearly resolve the majority of the Ddc2 peptide (residues 10-24), including the phosphoserine10 and 11 (**Figure 3A**). However, residues 4-9 were invisible due to flexibility and that they do not interact with the Rfa1-NTD. The Ddc2 peptide interacts in an analogous way to the *K. lactis* Ddc2 with ScRfa1-NTD. It forms a short α-helix and binds Rfa1-NTD OB-fold through a combination of hydrophobic and electrostatic interactions (**Figure 3B-C, Supplementary Figure 5D** and **E**). The conserved acidic residues of Ddc2 are complemented by a highly basic region of the protein-interaction cleft of Rfa1-NTD (**Figure 3C**). In our structure, similarly to those of *K. lactis* Ddc2-Rfa1-NTD, D14, D15, and D16 engage with the Rfa1-NTD, forming extensive salt-bridges with R44, K58, R93, and K95 (**Figure 3C**).

**Table 1.**
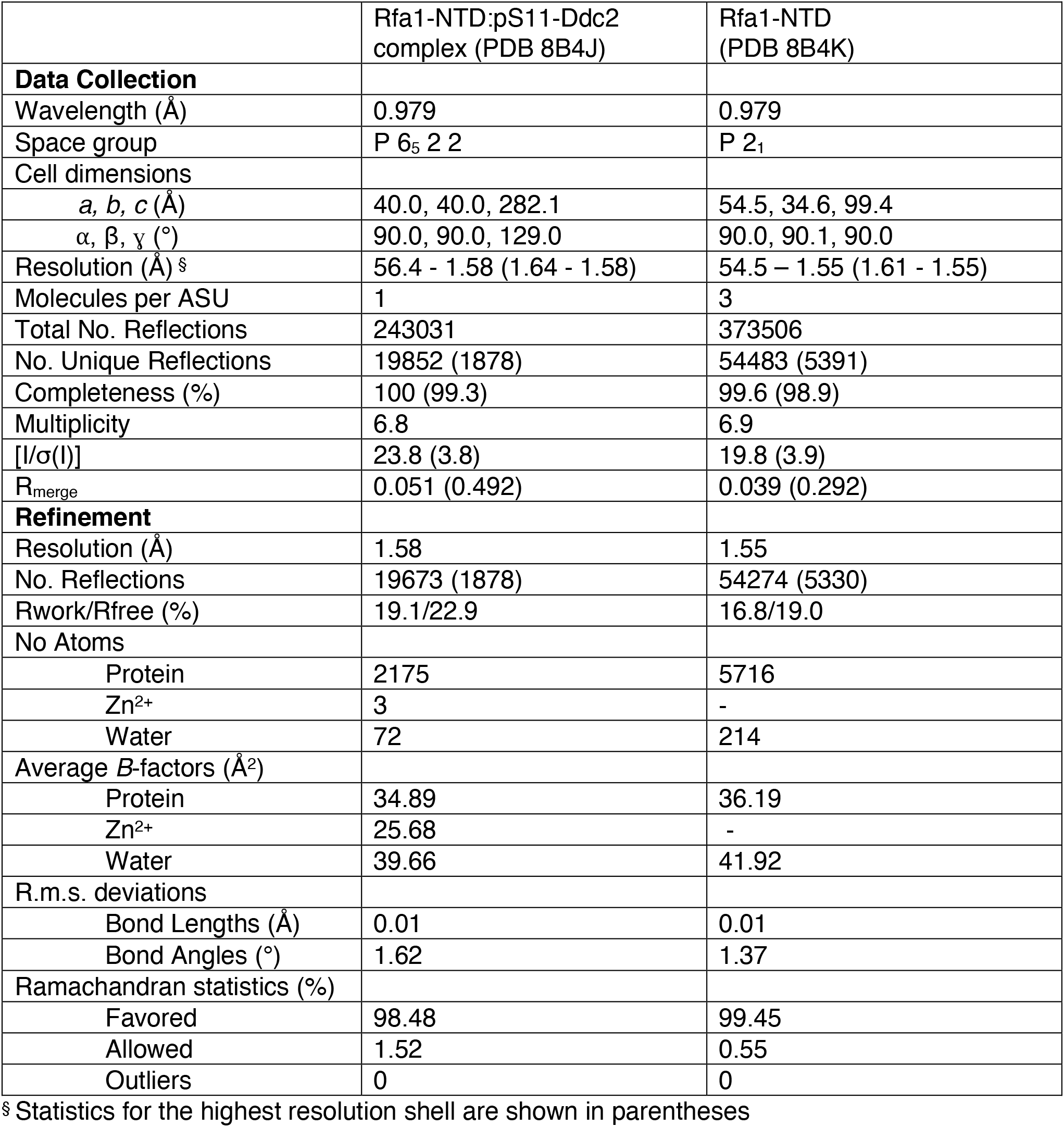
Crystallographic Data Collection and Refinement Statistics

**Figure 3.**
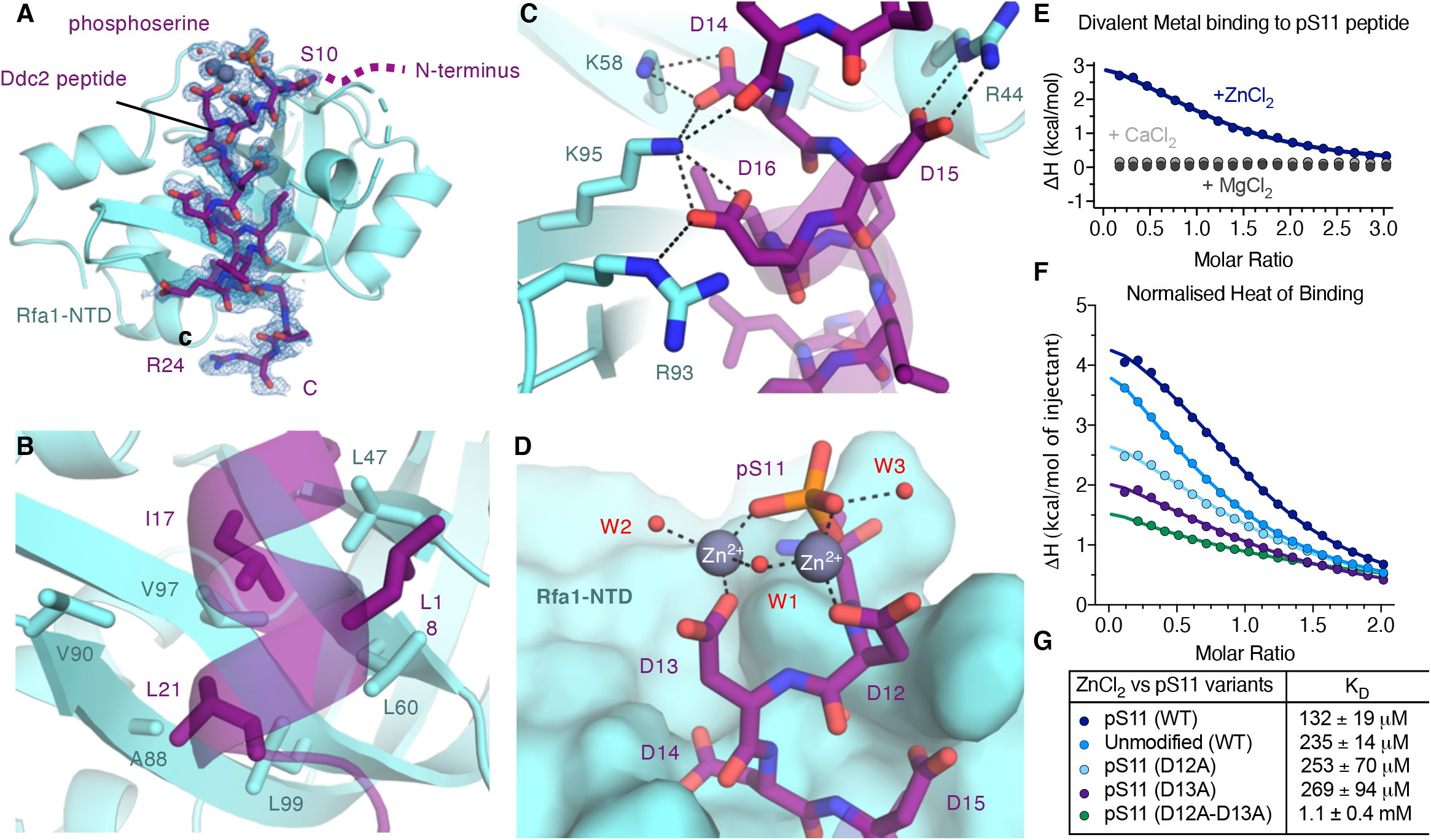
Structure of Rfa1-NTD in complex with pS11-Ddc2 peptide. **A** Crystal structure of Rfa1-NTD:pS11-Ddc2 peptide complex refined to 1.58 Å resolution. Rfa1-NTD is displayed as ribbon and rendered pale blue. Bound pS11-Ddc2 peptide is shown as sticks and rendered in purple. Heteroatoms are also coloured (red, oxygen; nitrogen, blue). Ordered waters and metal ions (Zn^2+^) are shown as red and grey spheres, respectively. Feature enhanced map (FEM) electron density, calculated in PHENIX, is shown as blue mesh (1 sigma) and carved around the peptide and metals (at 1.8 Å). **B-C** Molecular details of the binding site showing a combination of charge complementarity (b) and hydrophobic interactions (c). **D** Molecular details of Ddc2 peptide phosphoserine11 (pS11) coordinating two zinc ions (grey spheres), with 3 waters (red spheres) and Ddc2 residues D12 and D13. The RPA-NTD is shown as surface (pale blue) as this region of the peptide does not interact with the OB-fold. **E** Isothermal Titration Calorimetry (ITC) isotherm binding curves from metals titrated against pS11-Ddc2. Zinc ions (ZnCl_2_, bark blue) show clear binding, whereas both magnesium (MgCl_2_, dark grey) and calcium (CaCl_2_, light grey) show no detectable heat of binding. **F-G** ITC isotherm binding curves of ZnCl_2_ titrated against pS11-Ddc2 and calculated dissociation constants (KD). The binding affinity for Zn^2+^ decreases along with the heat of binding (ΔH, kcal/mol) when a zinc coordination donor is removed by substitution, with unphosphorylated, D12A, and D13A showing both a reduced affinity and heat of binding when compared to pS11-Ddc2 peptide. A double mutant D12A-D13A shows significantly diminished binding.

Critically, Asp12 and Asp13, together with phosphoserine11, coordinate two metal ions in the structure. The electron density from an omit map showed clear positive difference density for metal ions (**Supplementary Figure 6A**). Given that the crystallisation conditions contain ZnCl_2_, we postulate the metal ions to be Zn^2+^. Indeed, peptides that are aspartate-rich have been shown to bind to zinc ions (Miller et al., 2021), and in the structure, the ions are tetrahedrally coordinated by carboxyl groups of the aspartate side chains, oxygen groups of the phosphoserine, and water molecules (**Figure 3D**). The co-ordination geometry and bond lengths are consistent with zinc binding (∼2.0 Å in the structure) and were supported by using the CMM metal binding site validation server (Zheng et al., 2017). We further confirmed the pS11-Ddc2 peptide’s ability to coordinate zinc and no other divalent cations (magnesium or calcium) by ITC (**Figure 3E** and **Supplementary Figure 6B-D**). Clear binding was observed when ZnCl_2_ was titrated against the pS11-Ddc2 peptide, but not for either CaCl_2_ or MgCl_2_ (**Figure 3E** and **Supplementary Figure 6B-D**). To validate the zinc coordination observed in our structure, we performed additional ITC binding experiments using Ddc2 peptide variants. Our ITC data show that when a zinc coordination donor is removed by substitution, i.e. phosphoserine11→S11, D12A, or D13A, the apparent affinity decreases ∼2-fold compared to pS11-Ddc2 peptide (**Figure 3F and G**, and **Supplementary Figure 6E-H**). A double D12A/D13A substitution reduces the phosphopeptide affinity for zinc by ∼10-fold (**Figure 3F-G**). The importance of D12-D13 is underscored by its conservation in yeasts and humans, and is supported by earlier biochemical and genetic studies which show that D12K and D13K substitutions in Ddc2 results in defective RPA-Ddc2 interactions (Ball et al., 2007). To probe the Rfa1-NTD-pS11-peptide complex and zinc binding, we performed MST. As a control, the calculated dissociation constant for Zn^2+^ binding to the peptide was comparable to that obtained by ITC, and the affinity increases when the pS11 peptide is in complex with Rfa1-NTD but does not bind a range of other metals (**Supplementary Figure 7**). The zinc ion coordination in our structure is unusual, with phosphoserine and aspartate residues infrequently observed in zinc coordination sites in the Protein Data Bank (PDB) (Laitaoja et al., 2013). Generally structural zinc-coordination spheres are comprised predominantly of Cysteine and Histidine residues and possess nM-pM affinities for zinc (Laitaoja et al., 2013), with both Rfa1 and Mec1 both possessing such zinc sites (Tannous et al., 2021; Yates et al., 2018). The zinc coordination and the weak/moderate affinities measured, suggests that these metal ions are not likely to serve a structural role. Instead, pS11-Ddc2 may bind zinc with fluctuating metal concentrations in yeast; for example, during increased oxidative stress or when zinc is in excess.

### Ddc2 phosphorylation promotes oligomerisation of RPA

In the crystal lattice, we observe that the Ddc2 (pS11-D12-D13):Zn^2+^ coordinated complex, which protrudes from the Rfa1-NTD, contacts another Rfa1-NTD that completes the tetrahedral coordination of one of the zinc ions via H56 of the second Rfa1-NTD molecule (**Figure 4A and B**). This arrangement is reciprocated by the second Rfa1-NTD molecule forming a ‘fireman’s grip’ that maintains a 1:1 stoichiometry (**Figure 4A**), consistent with our ITC. Further, this second Rfa1-NTD (denoted a Rfa1-NTD’) contributes to the Ddc2 peptide binding via residues Y32, H56 and N122 **(Figure 4C)**. Specifically, Y32 contacts pS11 via a water molecule (**Figure 4B**). The zinc ion coordinated by Rfa1-NTD’-H56, Ddc2-D13 and Ddc2-pS11 is also stabilised by Rfa1-NTD’-F119 via cation-π interactions (**Figure 4B**). N122 also contacts the peptide backbone of Ddc2-D13. Structural analysis of this secondary interface by PISA (Krissinel and Henrick, 2007) suggests that it is stable in solution. We investigated this secondary site by MST using fluorescently labelled Ddc2 peptides and titrating Rfa1-NTD. This experimental set-up produces a biphasic binding curve by MST, indicating Ddc2 has a high-affinity and a low-affinity Rfa1-interaction site (**Figure 4C**). The first phase of binding occurs in the nanomolar to micromolar range, consistent with our ITC experiments, and the second phase in the micromolar range, which we attribute to the secondary binding site on Rfa1-NTD after the first site is saturated (**Figure 4C**). Focussed binding studies at Rfa1-NTD concentrations that have saturated the high-affinity site, we find that S11 phosphorylation enhances this interaction 3-fold (**Figure 4D**). The addition of zinc increases the affinity of Ddc2 for the secondary Rfa1-NTD further (∼7-fold) compared to the unmodified peptide (**Figure 4D**). This is consistent with our crystal structure, but also suggests that zinc is not obligatory for this interaction. We further investigated Rfa1-NTD dimer formation in solution by pS11-Ddc2 and zinc. We assessed the oligomeric state of Rfa1-NTD:pS11-Ddc2 peptide complexes using Analytical Size Exclusion Chromatography (**Figure 4F-H**). Our data showed that Rfa1-NTD exists as monomers and when in complex with Ddc2 peptide a peak shift is observed that is dependent on increasing concentrations of ZnCl_2_, which is more pronounced for pS11 peptide (**Figure 4F-H**). Cross-linking experiments provide further evidence for oligomerisation upon adding pS11 phosphopeptide and ZnCl_2_ and match the observations by SEC (**Supplementary Figure 8A**). Molecular weight estimations by SEC-MALS of Rfa1-NTD:pS11-Ddc2 peptide complex with ZnCl_2_ suggests a dimeric species compared to Rfa1-NTD alone (**Supplementary Figure 8B**). However, the broad peak suggests an equilibrium of dimers and monomers presumably caused by dilution effects during chromatography. The phosphorylation and zinc-enhanced dimerization of RPA also promotes RPA clustering on ssDNA as shown by MST experiments of RPA binding to ssDNA. The presence of pS11-Ddc2 peptide, without the coiled-coil domain, increases the hill coefficient of RPA binding to ssDNA from 1.2 to 1.5 and this is further increased to 1.8 when Zinc is added (**Supplementary Figure 8C-F**). These data are consistent with the idea that intra-RPA interactions, either via Rfa1-Rfa3 enhanced via Rfa1-S178 phosphorylation, binding to Ddc2-dimers, or phosphorylated Ddc2 (at S11) with zinc, promote cooperative clustering of RPA complexes on ssDNA.

**Figure 4.**
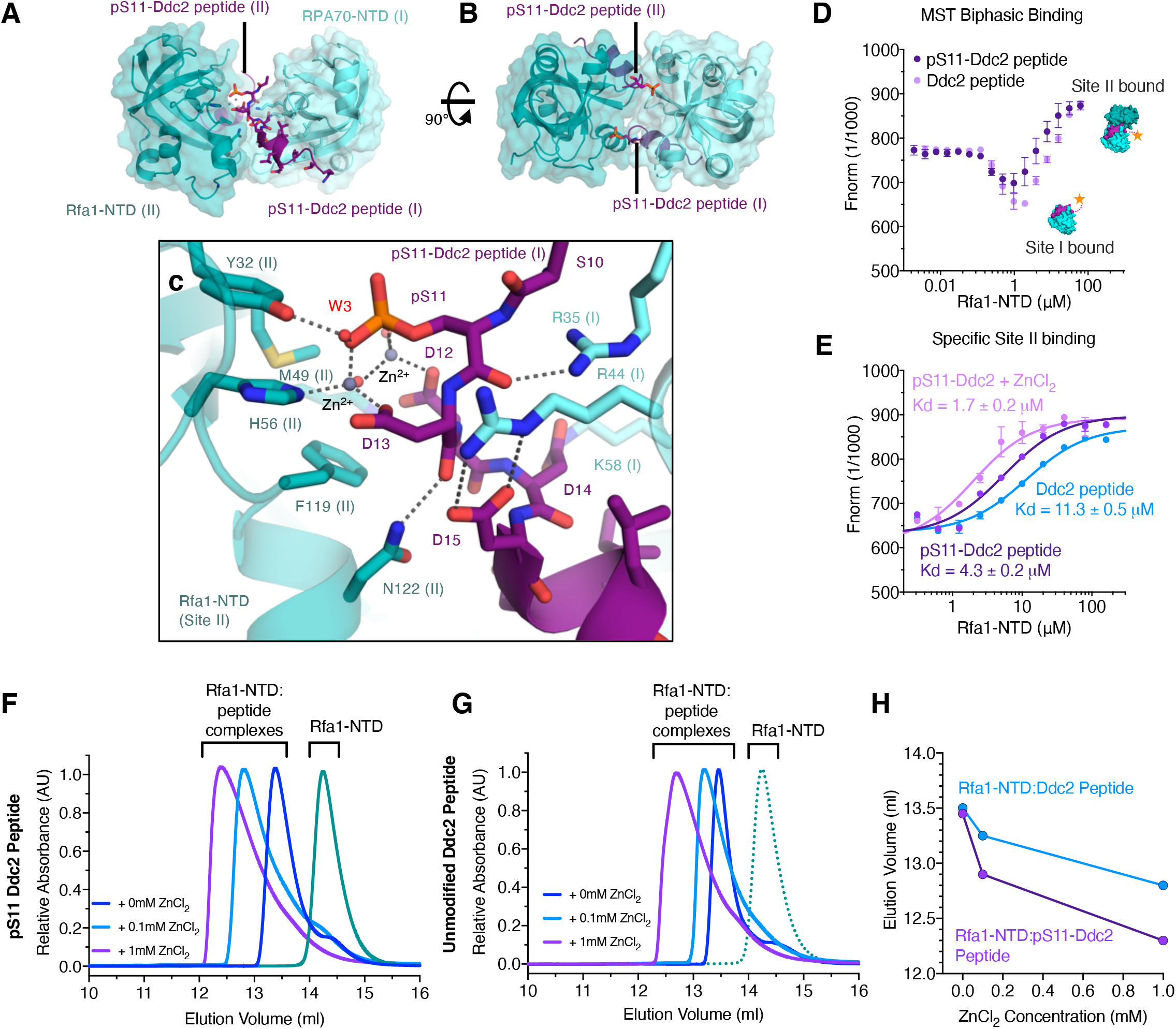
Rfa1-NTD dimer assembly via pS11-Ddc2 binding. **A-B** Orthogonal views of the Rfa1-NTD:pS11-Ddc2 complex dimer arrangement in the crystal lattice. Rfa1 and Ddc2 are coloured as in Figure 3. **C** Molecular details of the pS11-Ddc2 binding interface on the second Rfa1-NTD molecule. **D** Biphasic MST binding curves of 6-FAM labelled Ddc2 peptides versus Rfa1-NTD. **E** MST binding curves focussing on the secondary site, which occurs at Rfa1-NTD concentrations above site I saturation, with unmodified or S11 phosphorylated Ddc2 peptides and with 1mM ZnCl_2_. To obtain a Kd value MST curved were fitted with the law of mass action equation in Nanotemper analysis software. **F-G** size exclusion chromatography (SEC) of Rfa1-NTD in complex with (f) pS11-Ddc2 or (g) unmodified Ddc2 peptides and separated on a Superdex S75 (10/300) in buffer containing 0mM, 0.1mM and 1mM ZnCl_2_. **H** SEC peak elution volumes are plotted as a function of zinc concentrations.

Given the aspartate-rich nature of Ddc2 N-terminus, we considered acidophilic kinases that could target this motif – with Casein kinase II (CK2), which has the consensus motif S/T-X-X-D/E, being a plausible candidate. An analogous motif is found in *S. pombe* topoisomerase II has been shown to be a *bona fide* CK2 site (Nakazawa et al., 2019) (**Supplementary Figure 9A**). *In vitro*, we find that Ddc2-CC is a target of recombinant CK2, with two phosphorylation events observed upon kinase treatment (**Supplementary Figure 9B-C**). Substitution of both S10 and S11 to Alanine, but not individual serine substitutions, prevents phosphorylation (**Supplementary Figure 10D**). We thus used CK2 as a tool to investigate Ddc2-phosphorylation in vitro. Consistent with our peptide studies, phosphorylated Ddc2-CC, via CK2-treatment, shows a significant enhancement in binding to RPA compared to unphosphorylated protein (**Supplementary Figure 9E**). Further, size exclusion chromatography analysis shows higher-order oligomers of phosphorylated Ddc2-CC in complex with Rfa1-NTD upon addition of zinc (**Supplementary Figure 9F-G**). Superposing available structures, a multivalent arrangement of Rfa1-NTD:Ddc2-CC can be proposed, whereby the dimeric assembly of Rfa1-NTD:pS11-Ddc2 brings together and bridges two Ddc2-CC (thus Mec1-Ddc2) complexes (**Supplementary Figure 10**). To further investigate the potential higher order assembly, we visualised Mec1-Ddc2 complex using negative stain EM in the presence of Rfa1-NTD. We observe Mec1-Ddc2 complex particles (denoted here as monomers) as well as Mec1-Ddc2 clustering as dimers or higher order complexes that were closely linked in the presence of Rfa1-NTD (**Figure 5A-B**), a phenomena we had not seen with Mec1-Ddc2 alone (Sawicka et al., 2016; Tannous et al., 2021). The addition of zinc further promoted the clustering of Mec1-Ddc2 complexes (from ∼30% of total particles to ∼45%) (**Figure 5C**). Interestingly, using the recent advances in structure prediction by Alphafold (Jumper and Hassabis, 2022) in a Colab Alphafold multimer implementation (Mirdita et al., 2022) to ‘fill-in’ the missing coiled-coil (CC) and N-terminal portion of Ddc2 in our previously determined structure (Tannous et al., 2021), we produced a hybrid model of Mec1-Ddc2 in complex with the Rfa1-NTD (**Figure 5D**), which suggests that the kinase domains could face RPA, a known substrate, with the flexible linker allowing a range motion of the kinase. Combining these models with the Rfa1-NTD-Ddc2 dimer induced by phosphorylation, a supramolecular complex of two Mec1-Ddc2 molecules could form (**Figure 5E**), consistent with the observations in **Figure 5B-C**. Interestingly, several proteins involved in NHEJ are shown to act as bridges to bring two DNA-PK together (**Supplementary Figure 10**) (Chen et al., 2021), reminiscent of the higher order assemblies observed here.

**Figure 5.**
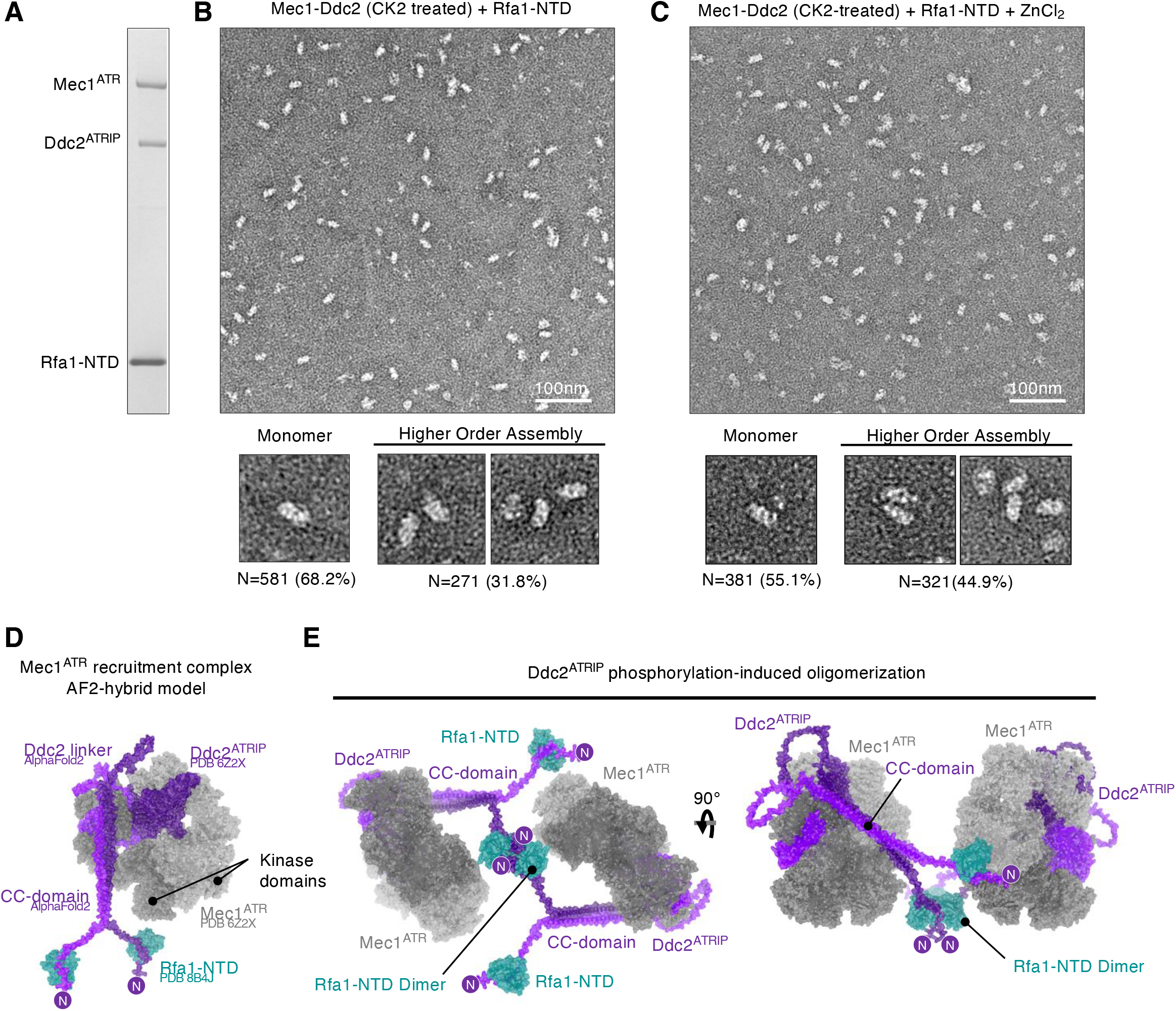
Model of Mec1-Ddc2 recruitment to RPA-coated ssDNA. **A** SDS-PAGE of the CK-treated Mec1-Ddc2 in complex with Rfa1-NTD. **B-C** Corresponding representative negative stain EM images in the absence (b) and presence of 0.1mM zinc (c). Individual particle images in large extraction boxes from each dataset are shown underneath describing the individual Mec1-Ddc2 particles (denoted monomers) and higher-order assemblies of closely associated complexes. The fraction of each type is provided from a semi-quantitative comparative analysis of the datasets. **D** A hybrid atomic model of Mec1-Ddc2 in complex with Rfa1-NTD was generated from Alphafold multimer predictions and experimentally determined structures of Mec1-Ddc2 complex (PDB 6Z2X) and Rfa1-NTD:pS11-Ddc2 peptide (determined in this study, PDB 8B4J). The Mec1 dimer is shown in shades of grey, and the Ddc2-dimer shown in shades of purple and the N-terminus of Ddc2 is highlighted. The Rfa1-NTD is coloured teal. **E** Orthogonal views of a supramolecular assembly of Mec1-Ddc2 complexes clustered through Rfa1-NTD:pS11-Ddc2 dimerization seen in our crystal structure.

## Discussion

Our study here combines biochemical, biophysical, crystallographic and yeast genetics studies, collectively supports a model where phosphorylation of RPA (Rfa1-S178) and Ddc2 (S11) promote the clustering of RPA on ssDNA and allow for further oligomerisation of Ddc2 via Rfa1-NTD. In turn, the oligomeric state of Mec1-Ddc2 acts to aid RPA clustering on ssDNA by tethering RPA complexes, and potentially enhancing recruitment efficiency. Specifically, our data show that RPA clustering stimulates Mec1-Ddc2 recruitment, which could lead to the phosphorylation of Rfa1 at S178 by Mec1 (Kim and Brill, 2003). The phosphorylation of S178 promotes RPA clustering, amplifies the recruitment of Mec1-Ddc2, and is consistent with Rfa1-S178A mutant RPA that has impaired Mec1 recruitment (Kim and Brill, 2003). Reciprocally, dimeric Ddc2 binding to RPA further enhances clustering of RPA through tethering. Studies on ATR have shown that localised crowding of ATR-ATRIP at the replication fork is important for the replication stress response (Yin et al., 2021) and the exchange of ATR-ATRIP at sites of RPA-coated ssDNA is critical for proper repair in murine models (Menolfi et al., 2018). The enhanced RPA clustering via phosphorylation of Rfa1-S178 and interactions with Ddc2, which are further promoted by Ddc2 (S11) phosphorylation and Zinc binding, creates a multi-faceted, intercalated and cooperative interaction network that can promote rapid accumulation (or removal) of Mec1-Ddc2. Mec1 recruitment dynamics may be an important aspect of regulating the checkpoint, particularly as the removal of RPA and Mec1-Ddc2 via Srs2 reduces Mec1-signalling (Dhingra et al., 2021).

Our study also demonstrates that phosphorylation of Ddc2, particularly at Serine11, enhances binding of Ddc2 to RPA. *In vivo and in vitro* studies performed here show that phosphorylation of Ddc2-S11 is important for the Mec1-checkpoint, but not its intrinsic kinase activity, consistent with its recruitment role. The structural and biophysical data suggest that Ddc2 phosphorylation of S11 also acts to promote further RPA clustering, through induced oligomerisation of the Rfa1-NTD, which can be further stabilised by Zn^2+^. In fact, immunoprecipitation experiments suggest that the RPA-interaction domain containing the N-terminal region of Ddc2 (residues 1-57) and lacking the dimeric CC domain promotes homodimerization of Ddc2 under DNA damaging conditions (Deshpande et al., 2017). In light of our structural and biophysical data, this could be interpreted as Ddc2 dimerization via Rfa1, in a phosphorylation-dependent manner. In addition, the structural arrangement observed here increases the overall binding surface area and strengthens the interaction. This may be important for localising Mec1-Ddc2 at short stretches of ssDNA where fewer number of RPA molecules are present. Greater numbers of RPA clustered on long ssDNA can clearly promote Chk1 phosphorylation by ATR-ATRIP (Choi et al., 2010), consistent with the model proposed here for RPA clustering and enhanced Mec1-Ddc2 recruitment, which could promote DDR foci formation. The flexible linker in Ddc2 would also allow a range of motion to enable Mec1 to phosphorylate many substrates, or other Mec1-Ddc2 complexes in our multivalent assembly. This arrangement will also likely promote Mec1 autophosphorylation, which occurs *in trans*, and is shown to be important for Mec1 function and repair efficiency (Hurst et al., 2021; Memisoglu et al., 2019; Sanford et al., 2021).

The role that zinc plays in the recruitment of Mec1 checkpoint kinase *in vivo* is less clear. In this study, we find that pS11-Ddc2 and the Rfa1-NTD:pS11-Ddc2 complex can specifically bind zinc ions over other metal ions, albeit with moderate/weak affinity. The presence of ZnCl_2_ improves complex formation of pS11-Ddc2:Rfa1-NTD, with the crystal structure only obtained when zinc was present. There are multiple examples of zinc ions at protein-protein interfaces that respond or are strengthened by transient increases in cellular zinc (reviewed in (Kocyła et al., 2021)). *Saccharomyces cerevisiae* has been shown to possess millimolar amounts of zinc stored in the vacuole (Simm et al., 2007), which can be transported inside the cell via vesicular compartments (Devirgiliis et al., 2004). In the cell, zinc is tightly bound in metal-protein complexes with limited labile pools of exchangeable ions. However, the overall available concentration of zinc more than exceeds the Kd values measured in our study, suggesting it is plausible that transient increases in cellular zinc, or zinc release through reductive stress (Li et al., 2022; Manford et al., 2021), may play a physiological role in the Mec1-checkpoint.

Both Mec1 and Ddc2 are extensively phosphorylated in response to DNA damage, replication stress, and during the cell cycle, in a Mec1-dependent and -independent manner, and are suggested to regulate the checkpoint (Bastos de Oliveira et al., 2015; Hustedt et al., 2015; Memisoglu et al., 2019; Paciotti et al., 2000). Two characterised Ddc2 phosphosites, S173 and S182, have been shown to permanently arrest the cell cycle in response to a DSB when mutated to an alanine (Memisoglu et al., 2019). Ddc2-S11 phosphorylation is suggested to be Cdk1-dependent *in vivo*, although S11 is not part of a canonical Cdk1 consensus motif (S/T-P-X-K/R, where X is any amino acid) (Holt et al., 2009). CDK activity is required for DSB processing and checkpoint maintenance via Rad9 (orthologue of human 53BP1) phosphorylation (Bonilla et al., 2008), so it is possible that this phosphosite is indirectly modified by Cdk1. *In vitro* we find that S10 and S11 can be phosphorylated by CK2, however we cannot rule out the activity of other Ser/Thr kinases involved in Ddc2-phosphorylation, such as Dbf4-dependent Cdc7 kinase (DDK). The exact kinase(s) responsible for phosphorylation of the Ddc2 N-terminal region will require more investigation but our study suggests that ancillary kinases operate to enhance Mec1/ATR checkpoint recruitment during replication stress or DNA damage response that could potentially be linked to cell cycle progression (Hanna et al., 1995).

## Methods

### DNA constructs

To generate the Rfa1-NTD construct, a DNA fragment encoding residues 1-132 was amplified using the RPA expression plasmid as a template and CloneAmp Hi-Fi PCR master mix (Clontech, Takara) following the manufacturer’s instructions. The DNA fragments were purified and cloned into a pOPINJ vector (a kind gift from Ray Owens, Addgene plasmid 26045) to encode a 3C-cleavable His_6_-GST-fusion using In-Fusion enzyme (Clontech, Takara), according to the manufacturer’s instructions. Ddc2-CC (residues 1-148) fragment was PCR amplified from yeast cDNA library with Phusion polymerase and digested with BamHI-XhoI (NEB) restriction endonuclease. The digested product was ligated into BamHI-XhoI digested pET28a (Novagen) using T4 ligase. The plasmid was sequence verified prior to its use in this study. To produce recombinant CK2 kinase the *CKA2* gene was PCR amplified from yeast cDNA library and cloned into a pOPINF vector (a kind gift from Ray Owens, Addgene plasmid 26042) to encode a 3C-cleavable His_6_ fusion protein. Single-stranded homopolymer oligonucleotides were purchased from IDT, as either 5′ Cy5 dT_100_ or unmodified dT_100_. For pull down assays 5′-Biotinylated-dT_100_ or 5’-Biotinylated-dT_32_ were purchased from IDT.

### Yeast strains, plasmids and proteins

Yeast strains used in this study were prepared as previously described (Tannous et al., 2021). In brief, PY434 (*MATa ade2-1 can1-100 his3-11,15 leu2-3,112 trp1-1 ura3-1 ddc2Δ::KanMX* containing pBL911 (pRS316, *URA3 DDC2*) was derived by genomic integration from C10-2A (*W303 RAD5 SML1*). Integration of *tel1Δ::NAT* in PY434 yielded PY436, and further integration of a *ddc1Δ::HIS3* cassette yielded strain PY437.The complementing plasmid pBL911 contains the *DDC2* gene on a centromeric plasmid under control of its own promoter, with *URA3* as both selectable and counter selectable (on 5-fluoroorotic acid, 5FOA) marker. Transformants were selected on YPD plates containing the respective drug and verified by PCR analysis. We then generated a centromere plasmid for each of the Ddc2 mutants p(*ddc2-x LEU2*) pBL912-x (pRS315, *ddc2-x LEU2*) for genetic analysis, and a 2-micron plasmid pBL904-x (pRS424, 2 µm ori *TRP1 MEC1 ddc2-x*) under control of the galactose-inducible *GAL1-10* promotor for overproduction of the Mec1-Ddc2-x complexes in yeast. All plasmids were verified by DNA sequencing.

### Purification of Ddc2 mutants

Strain PY439 (MATα *can1 his3 leu2 trp1 ura3 GAL pep4*∆:*:HIS3 nam7∆::KANMX4 ddc2∆::KANMX6 sml1∆::HYG*) was used to overexpress Mec1-Ddc2 or Mec1-ddc2-x mutants from the plasmid pBL904-x series, essentially as described previously (Tannous et al., 2021).

### Kinase assay

The kinase activity of Mec1-Ddc2 or Mec1-Ddc2-x complex was assessed as described previously (Tannous et al., 2021) using the kinase-dead version (K227A) of Rad53, fused to the Glutathione-S-transferase (GST) purification tag (GST-Rad53-kd), as a probe. The assay was performed in 10 µl of 25 mM Hepes-NaOH pH 7.4, 2 % glycerol, 1 mM DTT, 20 µg/ml BSA, 0.08 % ampholytes pH 3.5-10, 8 mM Mg-acetate, 100 µM ATP, 0.5 µCi [γ -^32^P] ATP and 100 mM NaCl, and Dpb11 activator as indicated.

### Yeast cell growth and analysis

Yeast strains PY434, PY436, and PY437 were transformed with plasmid pBL912 or its derivatives and selected on SC-Ura-Leu plates using standards yeast growth media. Transformants were grown overnight in SC-Leu medium and plated on SC-Leu plates containing 5-fluoroorotic acid (5FOA) to allow growth only of cells that had lost plasmid pBL911 (*URA3 DDC2*). These strains were grown in SC-Leu overnight at 30 °C and 10-fold serial dilutions spotted onto YPD plates, with or without hydroxyurea for varying times as indicated.

### Expression and Purification Mec1-Ddc2

Mec1-Ddc2 complexes, and its mutant variants, were expressed and purified essentially as described in ref (Tannous et al., 2021). A strain expressing Mec1 in complex with a N-terminal FLAG-tagged Ddc2 was a kind gift from J. Diffley. The yeast strain was grown at 30 °C in YP medium supplemented with 2% raffinose. When cultures reached 1–3 × 10^7^ cells/ml, expression was induced by the addition of 2% galactose for 2-3 h. Cells were harvested and washed in the lysis buffer (25mM Tris-HCl, 400mM KCl, 10% v/v Glycerol, 1mM EDTA, 0.01% NP40, 0.1% Tween-20, 1mM TCEP), then resuspended in lysis buffer supplemented with complete protease inhibitor cocktail tablets (Roche, 1 tablet per 50ml) at 0.3 x pellet volume, flash-frozen in liquid nitrogen and crushed using a 6875D Freezer/Mill Dual Chamber Cryogenic Grinderfreezer mill (SPEX SamplePrep). Cell powder from 12-18 l of yeast culture was resuspended in 25mM HEPES, 400mM KCl, 10% v/v Glycerol, 1mM EDTA, 0.01% NP40, 0.1% Tween-20, 1mM TCEP, supplemented with 2mM Benzamidine, protease inhibitor cocktail, and phosphatase inhibitors (5mM sodium pyrophosphate, 10mM Beta-glycerophosphate, 5mM sodium fluoride). The lysate was clarified by centrifugation at ∼20,000 x g for 30 minutes followed by ultracentrifugation at ∼200,000 x *g* for 60 min at 4 °C. Cleared lysates were incubated with pre-equilibrated Pierce™ anti-FLAG affinity resin (ThermoFisher) for 2 hours at 4°C with agitation. Beads were subsequently washed with 50 ml of 25mM HEPES, 400mM KCl, 10% (v/v) Glycerol, 1mM EDTA, 0.01% NP40, 0.1% Tween-20, 1mM TCEP, supplemented with protease inhibitor cocktail and phosphatase inhibitors, followed by incubation and washing with buffer containing ATP (5mM), and was finally washed with 50ml buffer. Protein was eluted with 25mM HEPES, 400mM KCl, 10% v/v Glycerol, 1mM EDTA, 0.01% NP40, 0.1% Tween-20, 1mM TCEP, 0.25-0.5mg/ml FLAG peptide. The eluates containing Mec1-Ddc2 were pooled and diluted to reduce the KCl concentration to ∼100mM and loaded onto a heparin column pre-equilibrated with 25mM HEPES, 100mM KCl, 10% v/v Glycerol, 1mM EDTA, 0.01% NP40, 0.1% Tween-20, 1mM TCEP and eluted using increasing concentrations of KCl to 400mM. Fractions containing good quality material, as assessed by SDS-PAGE, were pooled and used immediately without freezing.

### Expression and Purification of RPA

RPA was expressed and purified as described (Yates et al., 2018). Briefly, RPA expression vector, or mutants of, were transformed into BL21 (DE3) *E. coli*. A single colony from a plate was used to inoculate 1L of LB supplemented with kanamycin (34mg/L) and incubated overnight at 37°C without shaking. The culture was shaken at 170 rpm for several hours until an OD_600nm_ of 0.5-0.8. RPA expression was induced by the addition of 0.3mM (final) IPTG at 37°C for 3 hours. Cells were harvested by centrifugation at 5000x*g*. RPA was purified from the clarified lysate as described (Yates et al., 2018) using sequential affinity chromatography (Ni-NTA), anionic exchange (HiTrap Q), and gel filtration (Superdex S200 16/60). Purified RPA was assessed as >95% pure by SDS-PAGE and was nucleic acid free as assessed by its 260nm/280nm absorbance ratio (∼0.6) and was concentrated and buffer exchanged into 20mM HEPES, pH 7.6, 300mM NaCl, 1mM TCEP. Aliquots were flash frozen in liquid nitrogen and stored at -80°C until further use.

### Expression and Purification of Rfa1-NTD

His_6_-GST-Rfa1-NTD was expressed in BL21 (DE3) *E. coli*. This construct was constitutive overexpressed in *E. coli*, therefore a single colony from a transformant was inoculated per 1L of LB, supplemented with ampicillin (50mg/L), and incubated at 37°C without shaking overnight. The following morning the cultures were agitated at 180rpm until mid-logarithmic phase (OD_600nm_=0.5-0.8) and the temperature reduced to 20°C and incubated overnight with shaking. Cells were harvested by centrifugation at 5000x*g* and were frozen at -80°C. Cells were re-suspended in modified HEPES-buffered saline (HBS; 50mM HEPES, 150mM NaCl, 0.5mM TCEP, pH 7.5), lysed by sonication, and clarified by centrifugation. Protein was purified using its His_6_tag on Nickel Sepharose excel (cytiva). Once the His_6_-GST-fusion was adsorbed the Rfa1-NTD was liberated using on-column 3C cleavage overnight at 4°C and the resulting eluent purified further by size exclusion chromatography at 25°C in 20mM HEPES, pH 7.5, 150mM NaCl. The protein, which was assessed as >98% pure by SDS-PAGE, was concentrated and its concentration quantified spectroscopically using its theoretical extinction co-efficient and was either used immediately for crystallisation or flash frozen in liquid nitrogen for storage at -80°C. The protein is also stable at 4°C for several weeks.

### Expression and Purification of Ddc2-CC or GST-Ddc2-N

Ddc2-CC (residues 1-148) was expressed as N-terminally His_6-_tagged fusion protein in BL21 (DE3) *E. coli* using standard IPTG-induction procedures with an 18°C incubation overnight after induction. Cells were harvested by centrifugation and lysed by sonication. The protein was purified by NiNTA chromatography and eluted using increasing concentrations of imidazole, followed by ion exchange chromatography after dialysis into 50mM Tris, pH 7.5, 50mM NaCl, buffers, and polished using size exclusion chromatography in 20mM Tris, pH 7.5, 150mM NaCl using a Superdex 75 16/600. This construct contains no aromatic residues and therefore chromatographic steps were monitored by absorbance at 230nm. Protein concentration was quantified by BCA assay (Thermo Scientific). The protein was aliquoted, and flash frozen in liquid nitrogen stored in 20mM Tris, pH 7.5, 150mM NaCl. N-terminal fragments (and mutants thereof) of Ddc2 were expressed in BL21 (DE3) *E*.*coli* as GST-fusions using standard IPTG-induction procedures and 18°C incubation overnight after induction. Proteins were purified by glutathione Sepharose 4B (Sigma) as per manufacturer’s instructions using standard Tris-buffered saline buffers (20mM Tris, pH 7.5, 150mM NaCl). Protein concentration was quantified spectroscopically using its theoretical extinction co-efficient. The protein was aliquoted, and flash frozen in liquid nitrogen, and subsequently stored at -80°C until required.

### Synthetic Peptides

Synthetic Ddc2 peptides and phosphorylated and/or N-terminal 6-FAM labelled variants corresponding to residues 4-24 (sequence ETVGEFSSDDDDDILLELGTR) were commercially purchased from Pepmic Co. Ltd. (Suzhou, China) at >98% purity and milligram scale. Peptide sequences used in this study are indicated below (where superscript numbers indicate Ddc2 numbering, and [pS] indicates phosphoserine modification):

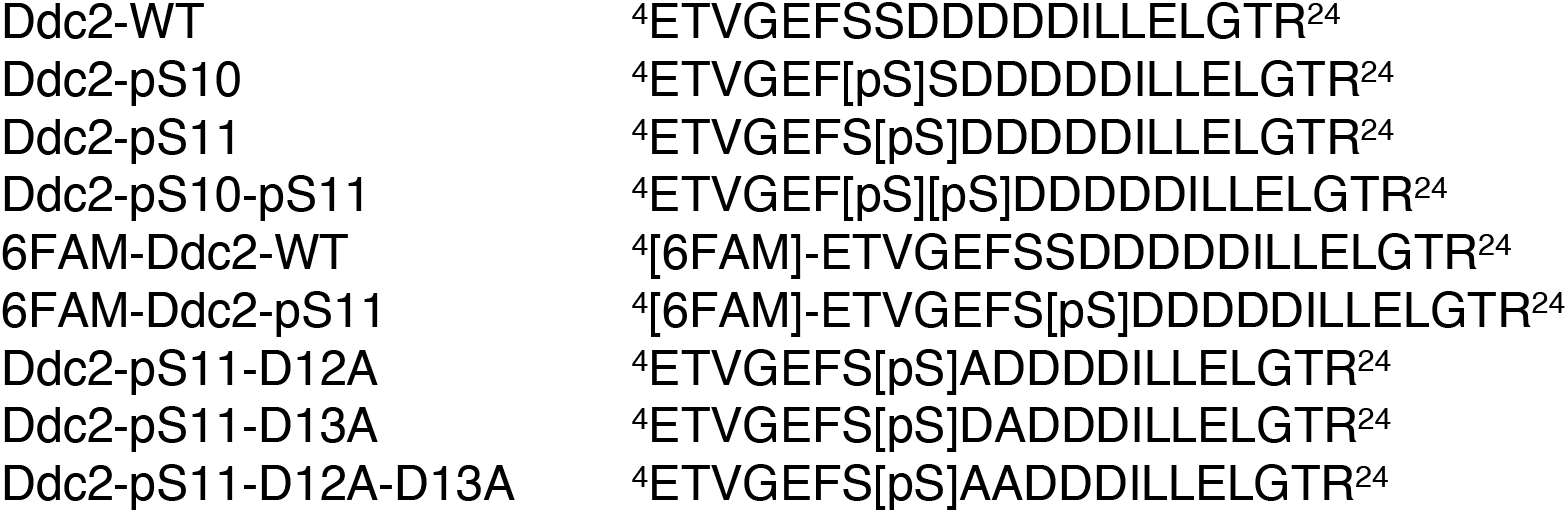

### CKA2

CK2 was expressed in BL21 (DE3) *E*.*coli* and expressed using standard IPTG-induction procedures and 18°C incubation overnight after induction. Cells were harvested by centrifugation and lysed by sonication. CK2 was purified using affinity chromatography (Nickel agarose), heparin-affinity chromatography, and gel filtration (Superdex 75 16/600) in 20mM Tris, pH 7.5, 150mM NaCl. Protein concentration was quantified spectroscopically using its theoretical extinction co-efficient. The protein was aliquoted, and flash frozen in liquid nitrogen, and subsequently stored at -80°C until required.

### Microscale Thermophoresis (MST)

MST experiments were performed using a Monolith NT.115 instrument (NanoTemper Technologies, Germany). In all cases premium treated capillaries were used and the experiment conducted at 25°C. Binding data was analysed using NanoTemper Analysis 1.2.101 software and plotted in GraphPad Prism. Each experiment was technically repeated at least three or more times and the mean half effective concentration (EC_50_), or Kd, values were calculated with Standard Error (SE) using the law of mass action equation.

### MST with RPA:Ddc2-CC vs ssDNA

RPA-ssDNA interaction studies used Cy5 5’ fluorophore labelled poly-thymine DNA oligos (5’Cy5-dT_100_). The fluorescently labelled DNA, in binding buffer (20mM HEPES, pH 7.6, 150mM NaCl, 0.5mM TCEP), was diluted to a concentration of 100nM and mixed with an equal volume of a serial dilution series of RPA or the phosphomimetic mutant RPA^S178D^ and incubated at room temperature for 20 minutes before loading into premium MST capillaries. Purified Ddc2-CC, or synthetic Ddc2 peptides, was added at to new dilution series experiments at a fixed concentration (25uM). MST experiments were performed using 80% LED power and 80% MST power with a wait time of 5 seconds, laser on time of 30 seconds and a back-diffusion time of 5 seconds. Each experiment was technically repeated at least three or more times and the mean half effective concentration (EC_50_) and Hill coefficient were calculated with Standard Error (SE).

### MST with Ddc2-CC vs RPA-ssDNA

RPA-ssDNA was generated on 5’Cy5-dT_100_ using a 4:1 M ratio of protein to oligonucleotide, based upon the previously determined 3-4 RPA molecules associating with dT_100_. The sample was diluted in binding buffer (20mM HEPES, pH 7.6, 150mM NaCl, 0.5mM TCEP) after a 20-minute incubation period room temperature to a final concentration of 200nM ssDNA, which is above the known Kd of RPA-ssDNA interaction. The RPA:Cy5-ssDNA complex was added to a serial dilution of purified Ddc2-CC in binding buffer incubated at room temperature for 20 minutes before loading into premium MST capillaries. MST experiments were performed using 80% LED power and 80% MST power with a wait time of 5 seconds, laser on time of 30 seconds and a back-diffusion time of 5 seconds. Binding data was fitted using NanoTemper Analysis 1.2.101. Each experiment was technically repeated at least three or more times and the apparent affinity (K_1/2_) were calculated with Standard Error (SE).

### MST with Phosphorylated Mec1-Ddc2 vs RPA-ssDNA

Purified Mec1-Ddc2 harbouring a 3xFLAG tag on the Ddc2 subunit was aliquoted into two samples. One half was treated with Lambda phosphatase and MnCl_2_ (NEB), the other half was treated with MnCl_2_ only, to produce a dephosphorylated sample and one that has its phosphorylation preserved. For MST experiments Mec1-Ddc2 was labelled by adding anti-FLAG antibody carrying an Alexa-647, on ice for 30 mins, at a 1:1M ratio. The subsequent sample was diluted to 100nM in 25mM HEPES, 100mM KCl, 10% v/v Glycerol, 1mM EDTA, 0.1% Tween-20, 1mM TCEP. The Mec1-Ddc2-mAb complex was added to a serial dilution of RPA-dT_100_ in 25mM HEPES, 100mM KCl, 10% v/v Glycerol, 1mM EDTA, 0.1% Tween-20, 1mM TCEP and incubated on ice for 20 minutes before loading into premium MST capillaries. MST experiments were performed using 80% LED power and 80% MST power with a wait time of 5 seconds, laser on time of 30 seconds and a back-diffusion time of 5 seconds. Binding data was fitted with the law of mass action equation using the NanoTemper Analysis 1.2.101 software. Each experiment was repeated at least three or more times.

### MST with phosphorylated Ddc2 peptides vs Rfa1-NTD

Synthetic Ddc2 peptides (residues 4-24, WT or pS11 variants) harbouring a 6-FAM were diluted in binding buffer (20mM HEPES, pH 7.6, 150mM NaCl) and used for MST experiments against a 2-fold serial dilution of purified Rfa1-NTD in binding buffer. The final concentration of fluorescently labelled peptide was 25nM. For metal interactions, Rfa1-NTD and 6-FAM pS11-peptide were incubated at 500nM in binding buffer at room temperature and used for MST experiments against a 2-fold serial dilution of metals diluted in binding buffer. The final concentration of fluorescently labelled peptide was 250nM. MST experiments were performed using 1-10% LED power and 80% MST power with a wait time of 5 seconds, laser on time of 30 seconds and a back-diffusion time of 5 seconds. Binding data was fitted with the law of mass action equation using the NanoTemper Analysis 1.2.101 software. Each experiment was repeated at least three or more times and at least twice for the metal interaction studies.

### EMSA

Binding assays were carried out in 20μl volumes. 5’Cy5-dT_100_ were mixed with purified RPA^WT^ or RPA^S178D^ at different molar ratios (1:1, 2:1, 3:1 protein:ssDNA) and diluted in binding buffer (20mM HEPES, pH 7.6, 150mM NaCl, 0.5mM TCEP) to a final ssDNA concentration of 200nM. Equimolar amounts of Ddc2 was added to as indicated. Nucleoprotein species were resolved by electrophoresis on an 3-12% PAGE run in 1X Tris-Acetate buffer at 60 V for 100 minutes at room temperature, loading 10ul of each reaction for each lane. The ssDNA complexes were visualized using in-gel fluorescence using a ChemiDoc gel imaging system (Bio-Rad). The experiment was performed several times with similar results.

### RPA-ssDNA capture assay

200nM 5’Cy5-dT_100_ and 5’-Biotinylated-dT_100_ were fully coated with purified RPA^WT^ or RPA^S178D^ by mixing with 4:1M ratio RPA to ssDNA, separately in 20mM HEPES, pH 7.6, 150mM NaCl, 0.5mM TCEP, 0.01% (v/v) Tween-20. Equal volumes of RPA-5’-Biotinylated-dT_100_ complexes were mixed with RPA-5’Cy5-dT_100_ complexes in the absence or presence of Ddc2-CC (200nM). Each reaction (50ul volumes) were incubated with 10μl pre-equilibrated magnetic streptavidin resin and incubated for 15 minutes at room temperature in the dark. The resin was pulled down using a magnet and washed 2 times with 50 ul of 20mM HEPES, pH 7.6, 150mM NaCl, 0.5mM TCEP, 0.01% (v/v) Tween-20. The beads were resuspended in 1X NuPAGE LDS loading buffer and the beads boiled at 95 °C for 5 minutes in a dry heat block. The resin was separated from the samples using a magnetic rack, which were loaded onto a 4-12% Bis-Tris gradient SDS-PAGE (ThermoFisher) and run in 1X MES running buffer (ThermoFisher) for 30 minutes. The ssDNA was visualized using in-gel fluorescence using a ChemiDoc gel imaging system (Bio-Rad) and the protein stained using ReadyBlue protein stain (Sigma-Aldrich). Fluorescent DNA bands and protein bands were quantified using the Bio-Rad Image Lab software 6.0.1.

### Mec1-Ddc2: RPA Pull-down

5’ Biotinylated-dT_32_ was immobilized onto pre-equilibrated 20μl streptavidin coated magnetic beads (Thermo Scientific) at saturating concentrations before being washed with several 1ml volumes of wash buffer (50mM Tris-HCl, pH 7.5, 150mM NaCl) and the beads separated using a magnet. The ssDNA-conjugated beads were incubated with RPA^WT^ or RPA^S178D^ for 15 minutes at room temperature before being washed as before. Purified Mec1-Ddc2 was incubated with the immobilized RPA-dT_32_ complexes for 30 minutes on ice. Mec1-Ddc2-RPA-dT_32_ complexes were separated using a magnet and the beads re-suspended and washed three times wash buffer. Finally, the beads were separated using a magnet and mixed with equal volumes of NuPAGE LDS-loading buffer (ThermoFisher) before the proteins were resolved by SDS-PAGE. Binding control experiments were performed with strepdavidin-5’Btn-dT_32_ in the absence of RPA and RPA-dT_32_ in the absence of Mec1-Ddc2.

### Gel filtration analysis

Rfa1-NTD in complex with near equimolar amounts of either Ddc2-peptide or pS11 Ddc2-peptide (200 μl samples) were injected at a concentration of 10mg/ml onto a Superdex S75 (10/300, Cytiva) in 25 mM HEPES, pH 7.0, 100 mM NaCl, with ZnCl_2_ at indicated concentrations, at 25 °C, with a flow rate of 0.4 ml/min. The gel filtration column was equilibrated in the appropriate buffer between runs.

### SEC-MALS

Rfa1-NTD in complex with near equimolar amounts of pS11 Ddc2-peptide (200 μl) was injected at a concentration of 10mg/ml onto a Superdex S75 (10/300, Cytiva) mounted on a high-pressure liquid chromatography system (1260 Infinity; Agilent). Species were separated using 25 mM HEPES, pH 7.0, 100 mM NaCl, 1 mM ZnCl_2_, at 25 °C, with a flow rate of 0.4 ml/min. Real-time light scattering and refractive index were simultaneously measured (Helios-II, T-rEX; Wyatt). Rfa1-NTD was also run alone as a control to compare relative MW estimates. The Astra software package was used for data analysis (Wyatt).

### Cross-linking SDS-PAGE

Reactions were carried out in 20μl volumes. Rfa1-NTD in complex with near equimolar amounts of either Ddc2-peptide or pS11 Ddc2-peptide in 25 mM HEPES, pH 7.0, 100 mM NaCl, with ZnCl_2_ at indicated concentrations. Glutaraldehyde cross-linker (0.05% final concentration) was added and the sample incubated for 5 minutes at 25 °C. A no cross-linker was also included. The samples were quenched using a 1x NuPAGE LDS loading buffer supplemented with 100mM Tris-HCl, pH 7.5. Samples were boiled at >95°C for 5 minutes in a dry heat block before being resolved by SDS-PAGE.

### Isothermal Titration Calorimetry

ITC measurements were performed on a MicroCal PEAQ-ITC calorimeter (Malvern Panalytical). Data were analyzed using the MicroCal PEAQ-ITC analysis software supplied by the manufacturer using the One Set of Sites model. Titrations were carried out at 750 rpm and at 25°C. For each titration, the heat change associated with ligand dilution was measured and subtracted from the raw data.

### Rfa1-NTD vs phosphopeptides

Commercially purchased synthetic Ddc2 peptides (and phosphorylated variants) were dissolved in 25mM HEPES, pH 7.0, 100mM NaCl, using a volume to produce a stock concentration of 1mM. Rfa1-NTD, or peptide solutions (0.5mL volumes) were dialyzed overnight at 4°C against 1L of 25mM HEPES, pH 7.0, 100mM NaCl, using a 1kDa cut-off dialysis cassette (Cytiva), with agitation with magnetic stirrer. Protein concentration was determined by absorbance at 280nm using a UV-2550 spectrophotometer. Peptide concentrations were adjusted based on the final volumes obtained after dialysis. Samples were diluted to a working concentration in 25mM HEPES, pH 7.0, 100mM NaCl, 5% (v/v) glycerol and centrifuged at 13,000 rpm for 15 minutes at 4°C to remove aggregates prior to ITC measurements. The peptide or phosphopeptides were loaded in the syringe at a concentration of 475μM, and Rfa1-NTD placed into the cell at a concentration of 50μM, or buffer (25mM HEPES, pH 7.0, 100mM NaCl, 5% (v/v) glycerol) for heat of dilution controls.

### Zinc interaction studies

For metal-peptide interaction studies ZnCl_2_, MgCl_2_, and CaCl_2_ stock solutions were prepared at a working concentration of 5mM in 25mM HEPES, pH 7.0, 100mM NaCl and filtered through a 0.22μm filter and loaded into the syringe. Ddc2 peptides (and phosphorylated variants) dialysed and diluted in 25mM HEPES, pH 7.0, 100mM NaCl, at a concentration of 500 μM, were placed in the cell for interactions. Buffer (25mM HEPES, pH 7.0, 100mM NaCl) was placed in the cell for heat of dilution controls.

### Crystallisation and Structure Determination

Lyophilised commercially bought synthetic pS11-Ddc2 peptide (^4^ETVGEFS[pS]DDDDDILLELGTR^24^, where [pS] refers to a phosphoserine) was dissolved in purified and concentrated Rfa1-NTD (20 mg/ml in 20mM HEPES, 100mM NaCl, 0.5mM TCEP) so that the phosphopeptide: protein molar ratio was 3:1. The resulting phosphopeptide protein complex was centrifuged at 13,000 rpm at room temperature to remove undissolved solids. The complex was crystallised in; 10 mM Zinc Chloride, 0.1M MES pH 6.0, 20% w/v PEG 6000 at 22°C using nanolitre sitting drop vapour diffusion. Crystals formed from heavy precipitation and grew over 2 weeks via Ostwald ripening. Rfa1-NTD was also crystallised alone in 0.2 M lithium chloride, 0.1 M sodium acetate, pH 5.0, 20% (w/v) PEG 6000. Crystals were cryo-protected by the addition of ethylene glycol to a maximum concentration of 25% (v/v) before mounting and flash-freezing in liquid nitrogen. Diffraction data were collected on beamline I03 and I04 at Diamond Light Source, Didcot, Oxfordshire, UK, using an Eiger2 XE 16M detector under cryogenic conditions. Diffraction revealed two crystal forms of space groups *P*6_5_22, for the phosphopeptide complex, and *P*2_1_, for Rfa1-NTD alone, with unit cell parameters of *a* = 40.0 Å, *b* = 40.0 Å, *c* =282.1 Å, *α* = 90.0°, *β* = 90.0°, *γ* = 129°, and *a* = 54.5, *b* = 34.6, *c* = 99.4 Å, *α* = 90.0°, *β* = 90.1°, *γ* = 90.0°, for the Rfa1-NTD alone. Data were auto-processed using Xia2-DIALS (Winter et al., 2022) routines during data collection at the beamline. Data was reduced in CCP4 package (Winn et al., 2011) to produce the best quality data resulting in two complete (>99% in the highest resolution shell) datasets with resolutions of 1.58 and 1.55 Å for the *P*6_5_22 and *P*2_1_ crystal forms, respectively, and an overall *R*_merge_ ∼5% for both datasets. The crystal structures were determined by molecular replacement using Phaser and the PDB 5OMB. One molecule of Rfa1-NTD was found per asymmetric unit (ASU) for the *P*6_5_22 space group, and 3 molecules per ASU for the *P*2_1_ crystal form. Clear positive difference density corresponding to the pS11-Ddc2 peptide was observed after molecular replacement and was manually built in Coot (Emsley and Cowtan, 2004). Structures were refined in the PHENIX package (Adams et al., 2010; Liebschner et al., 2019) and subsequent models adjusted and rebuilt in Coot. During refinement rounds, riding hydrogens were added to the model, and TLS restraints were also applied. The final structures possessed good R-factors as well as excellent model geometry (see Table 1). Both structures possessed zero Ramachandran outliers with 100% of the residues within allowed regions (>98% in favoured regions), as assessed by MolProbity (Chen et al., 2015). Models and structure factors have been deposited in the RCSB PDB with accession codes 8B4J (*P6*_*5*_*22*_1_, 1.58 Å, Rfa1-NTD:pS11-Ddc2 complex) and 8B4K (*P*2_1_, 1.55 Å, Apo). Structural figures were prepared using PyMol (Schrödinger).

### PhosTag Ddc2 phosphorylation assay

Reactions were carried out in 20μl volumes. Purified Ddc2-CC (50 µM ∼1mg/ml) was mixed with a serial dilution of purified CK2α’ indicated (6μM – 6nM) in Tris Buffered Saline (Tris-HCl, pH 7.5, 150mM NaCl, 0.5mM TCEP) supplemented with 5mM ATP and 10mM MgCl_2_ and incubated at 25°C. The reaction was stopped by the addition of SDS-PAGE sample buffer and boiled at >95°C for 5 minutes in a dry heat block. To determine phosphorylation over time a large kinase master mix was prepared using purified Ddc2-CC (50 μM, ∼1mg/ml) and 1 μM CK2α’. Reactions were initiated by the addition of 5mM ATP with 10mM MgCl_2_ (final concentrations), and 20μl samples taken and stopped by the addition of SDS-PAGE sample buffer and boiled at >95°C for 5 minutes in a dry heat block at indicated time points after mixing and incubating at 25°C. Samples were either resolved by Tris-Glycine SDS-PAGE, or SuperSep precast gels containing 50μM Phos-tag™ Acrylamide (Fujifilm), which shifts protein bands based on phosphorylation status. Gels were run according to manufacturer’s instructions.

### Negative Stain EM

Mec1-Ddc2 complexes expressed and purified essentially as described (Tannous et al., 2021), were treated with 100nM recombinant CK2 and 1mM ATP with 4mM Mg(OAc)_2_ for 1 hour at room temperature. The sample was diluted to ∼170nM, and equimolar amounts of Rfa1-NTD were added. The sample was split in half and ZnCl_2_ was added (100μM final concentration), or buffer without zinc. 4μl from each sample was applied to freshly glow-discharged carbon-coated grids and incubated for 1 minute. The excess sample was blotted, and two 4 μl washes of distilled water were applied. The sample was stained by applying 4 μl of 2% (w/v) filtered uranyl acetate for 30s and the excess blotted and allowed to air dry. Grids were imaged on a FEI Tecnai G2 spirit 120kV electron microscope equipped with a 2K camera at a nominal magnification of 67k, resulting in a pixel size of 3.5 Å. All images were collected manually with a defocus range from -2.5 to -3.5 μm. A semi-quantitive approach was taken to ‘count’ Mec1-Ddc2 oliomers. Particles were picked from 10 micrographs using gautomatch, and were extracted and classified in Relion 3.0 (Zivanov et al., 2018) using a box size of 256 pixels (896 Å) but a small circular mask of 300 Å. Particles that presented clear expected features of Mec1-Ddc2 were kept and classified further to remove junk. The raw extracted particle images from ∼900 particles of each dataset were used to assess the number of higher order oligomers. These were defined as particles within 100 Å (the length of the Ddc2-CC domain) of each other.

## Contributions

L.A.Y., and X.Z. designed the project. Protein construct cloning, purification, and biophysical studies were performed by L.A.Y, and yeast genetic studies were carried out by E.A.T. R.M.M., collected X-ray data. L.A.Y., carried out the structural studies. L.A.Y. and X.Z. analysed the results and wrote the manuscript with contributions from E.A.T., and P.J.B. All authors assisted with manuscript editing.

## Acknowledgments

X-ray data collection was done at Diamond light source (Oxfordshire, UK). We thank Diamond for beamtime (proposal mx23620) and the staff of beamlines I03 and 104. We thank the Centre of Structural Biology (CSB) for crystallization and EM support. We thank Dr L. Masino and the biophysics technology platform (Francis Crick Institute), for their support with ITC data, and Zhang and Burgers laboratory members for helpful discussions. This work was funded in part by Wellcome Trust grant no. 210658/Z/18/Z (to X.Z.) and by grant no. GM118129 from the National Institutes of Health (to P.M.B.).

## Competing Interests

The authors declare no competing interests.

## Notes

### Competing Interest Statement

The authors have declared no competing interest.

