## Supplementary Figure for "A DNA damage-induced phosphorylation circuit enhances Mec1^ATR^-Ddc2^ATRIP^ recruitment to Replication Protein A"

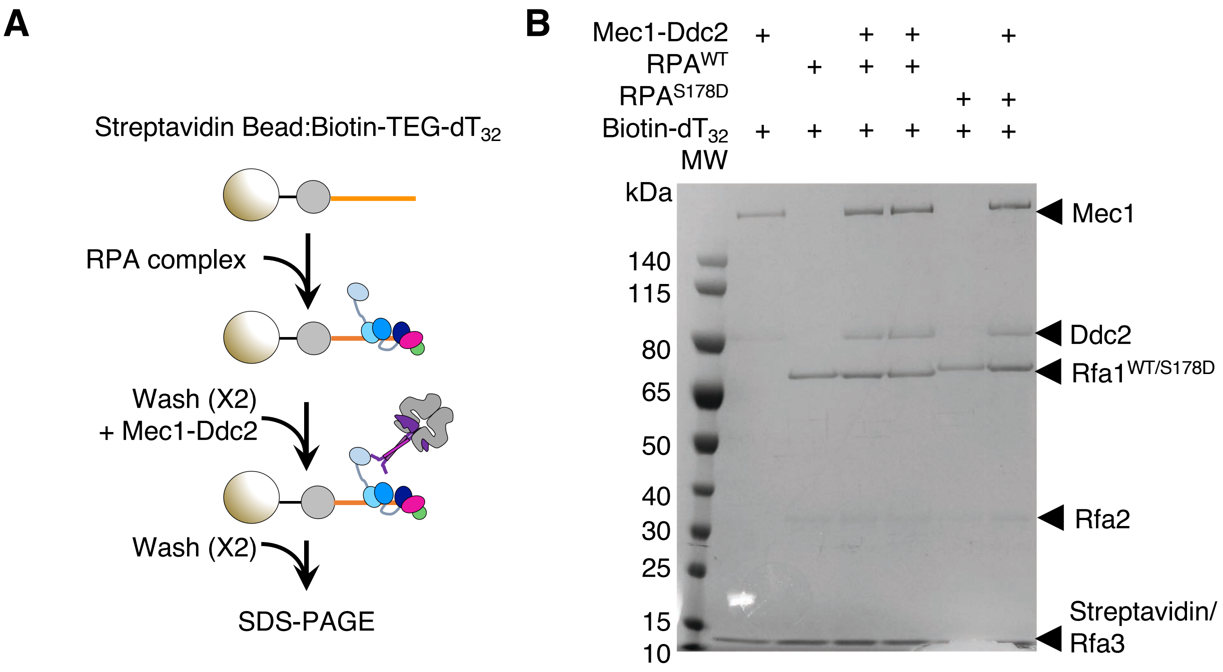


**Supplementary Figure 1**

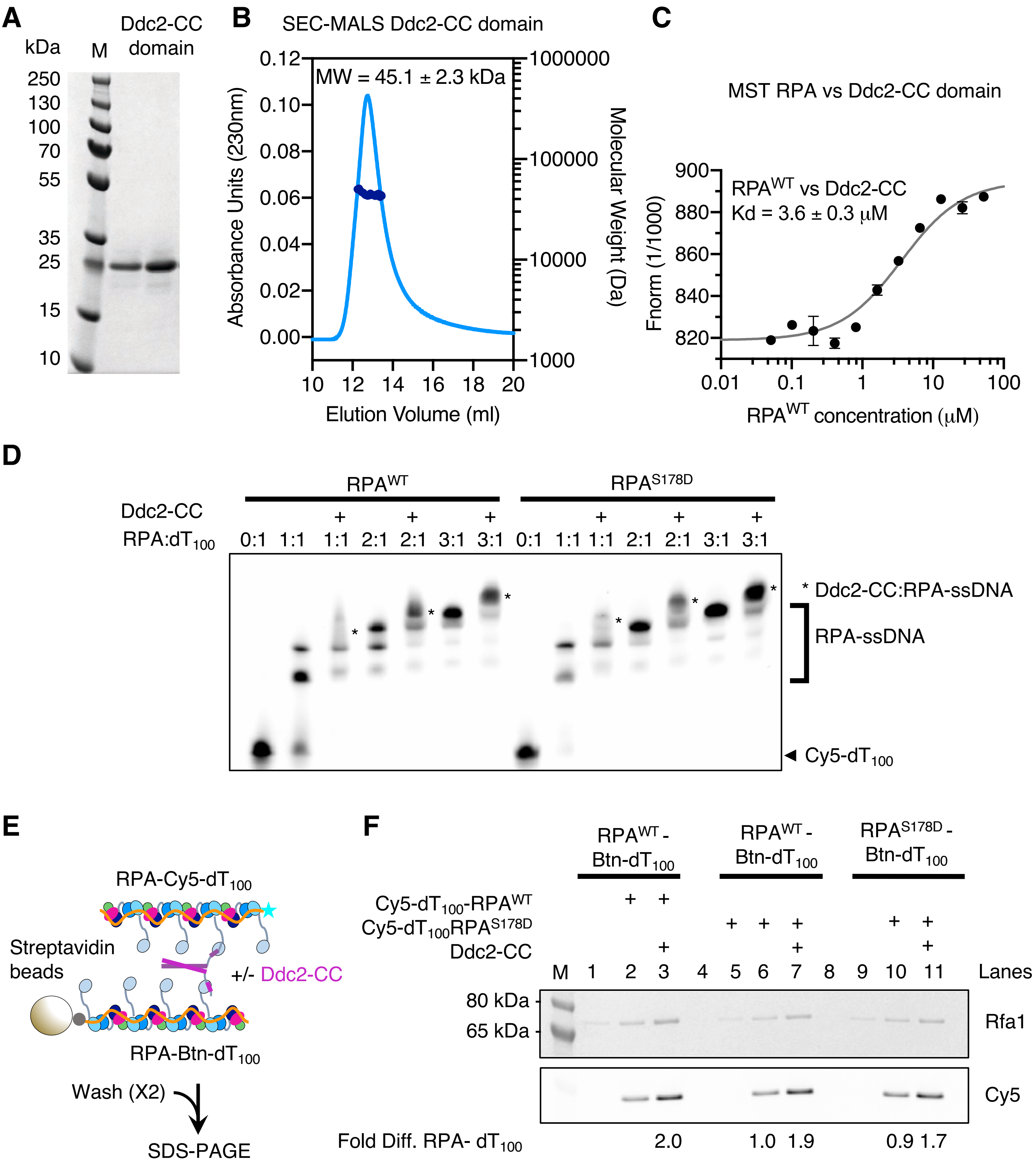


**Supplementary Figure 2**

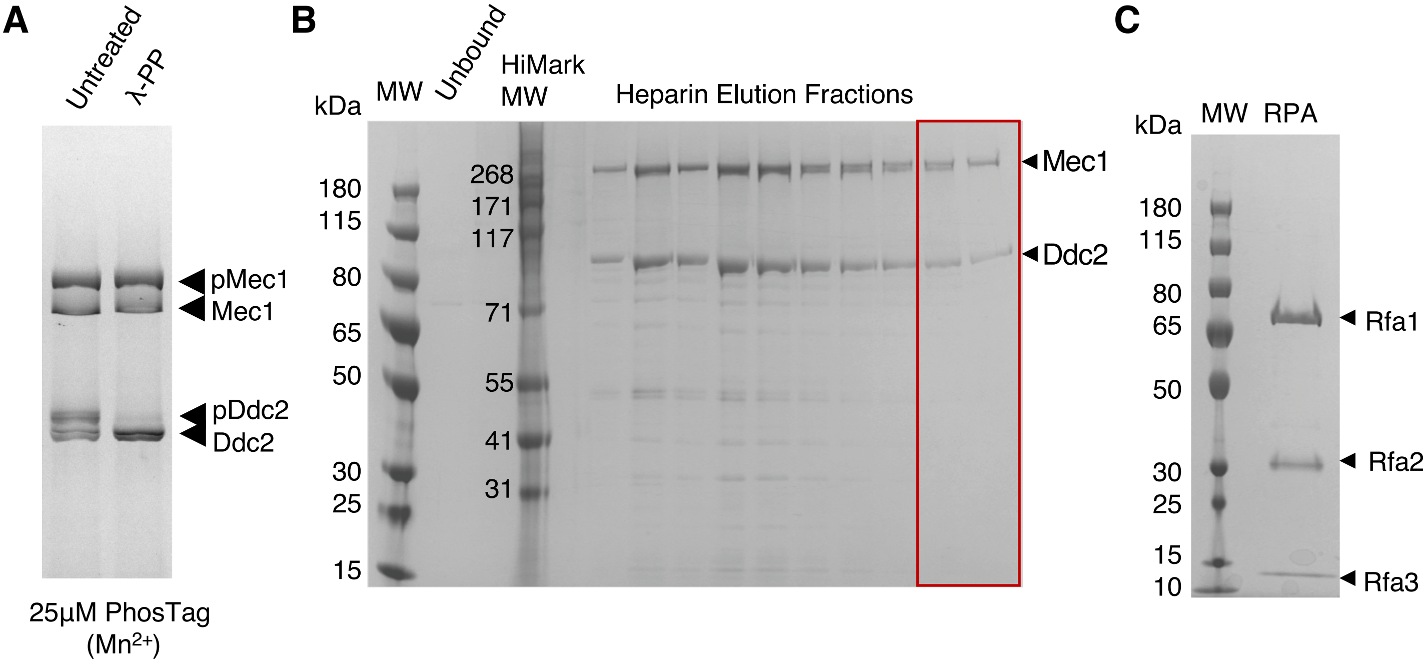


**Supplementary Figure 3**

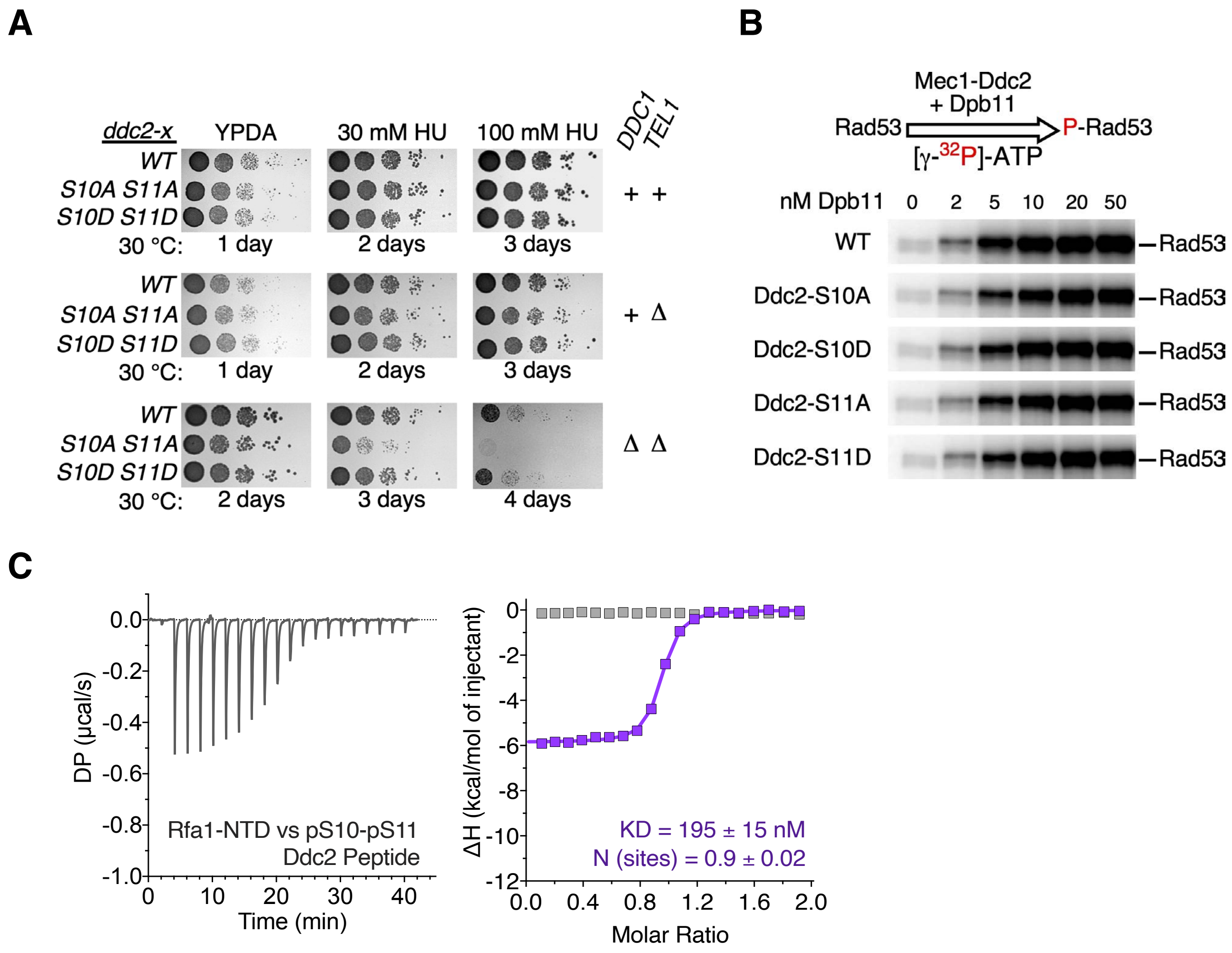


**Supplementary Figure 4**

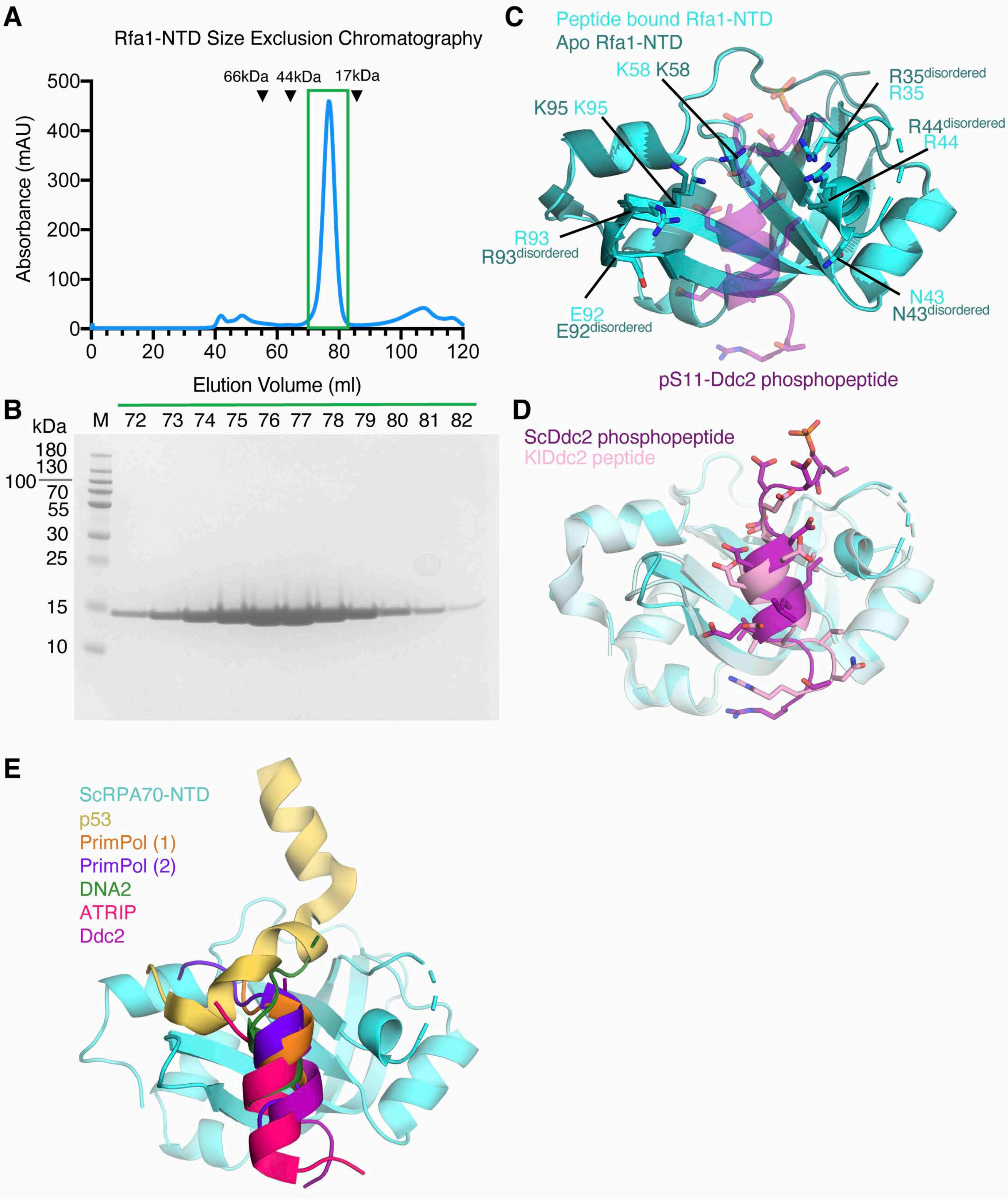


**Supplementary Figure 5 – Rfa1-NTD purification and structure comparisons**.

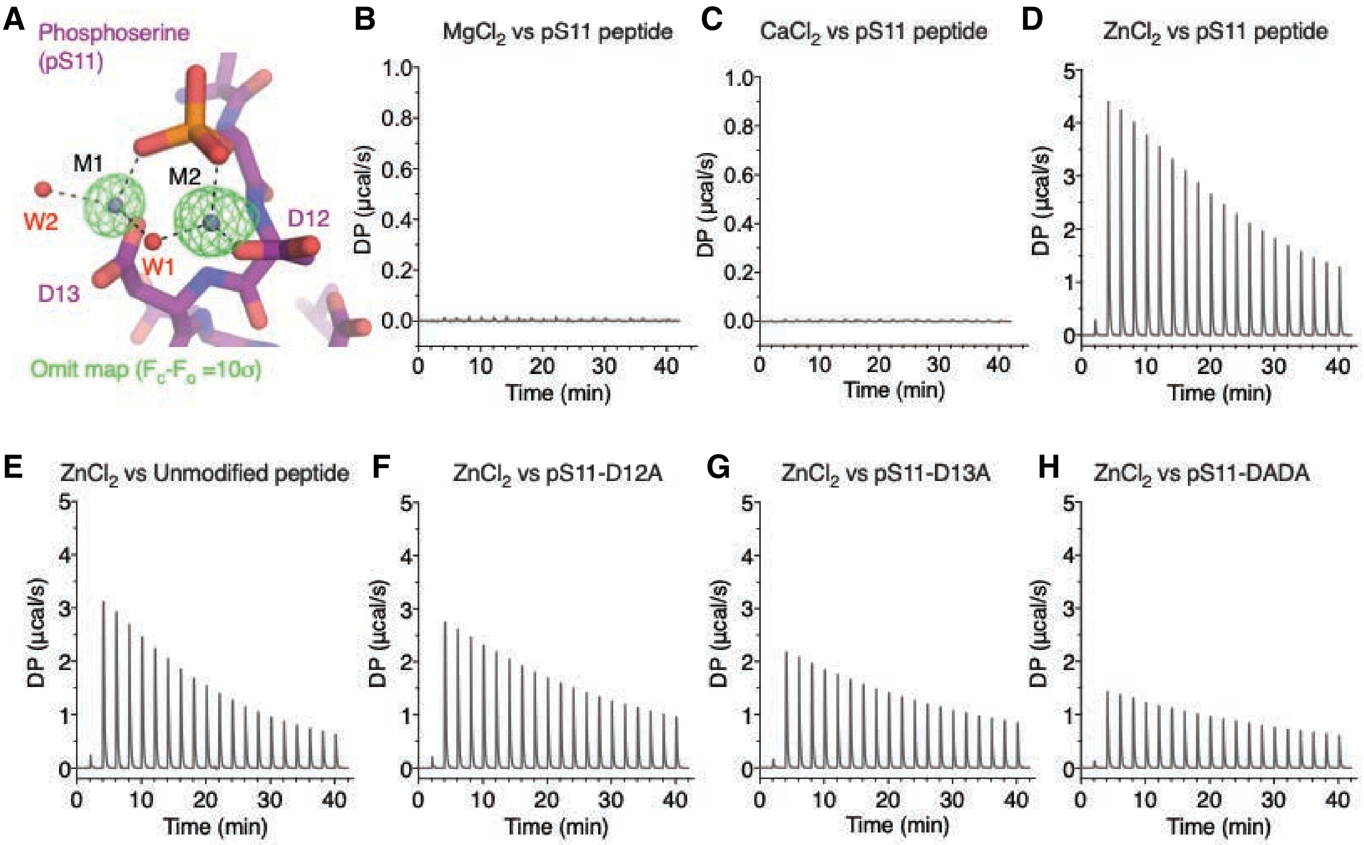


**Supplementary Figure 6**

**A**

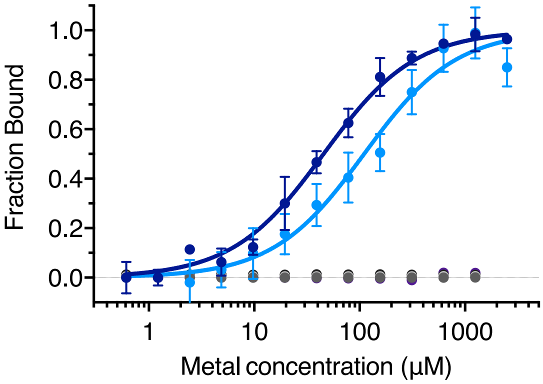


**B**

| Metal Interactions | Kd (μM) |
|---|---|
| ● pS11-Ddc2 Peptide vs $ZnCl_2$ | 110.3 ± 13 |
| ● Rfa1-NTD:pS11-Ddc2 complex vs $ZnCl_2$ | 37.7 ± 5 |
| ● Rfa1-NTD:pS11-Ddc2 complex vs $CaCl_2$ | n.d. |
| ● Rfa1-NTD:pS11-Ddc2 complex vs $MgCl_2$ | n.d. |
| ● Rfa1-NTD:pS11-Ddc2 complex vs $MnCl_2$ | n.d. |
| ● Rfa1-NTD:pS11-Ddc2 complex vs Fe(II)$SO_2$ | n.d. |

**Supplementary Figure 7**

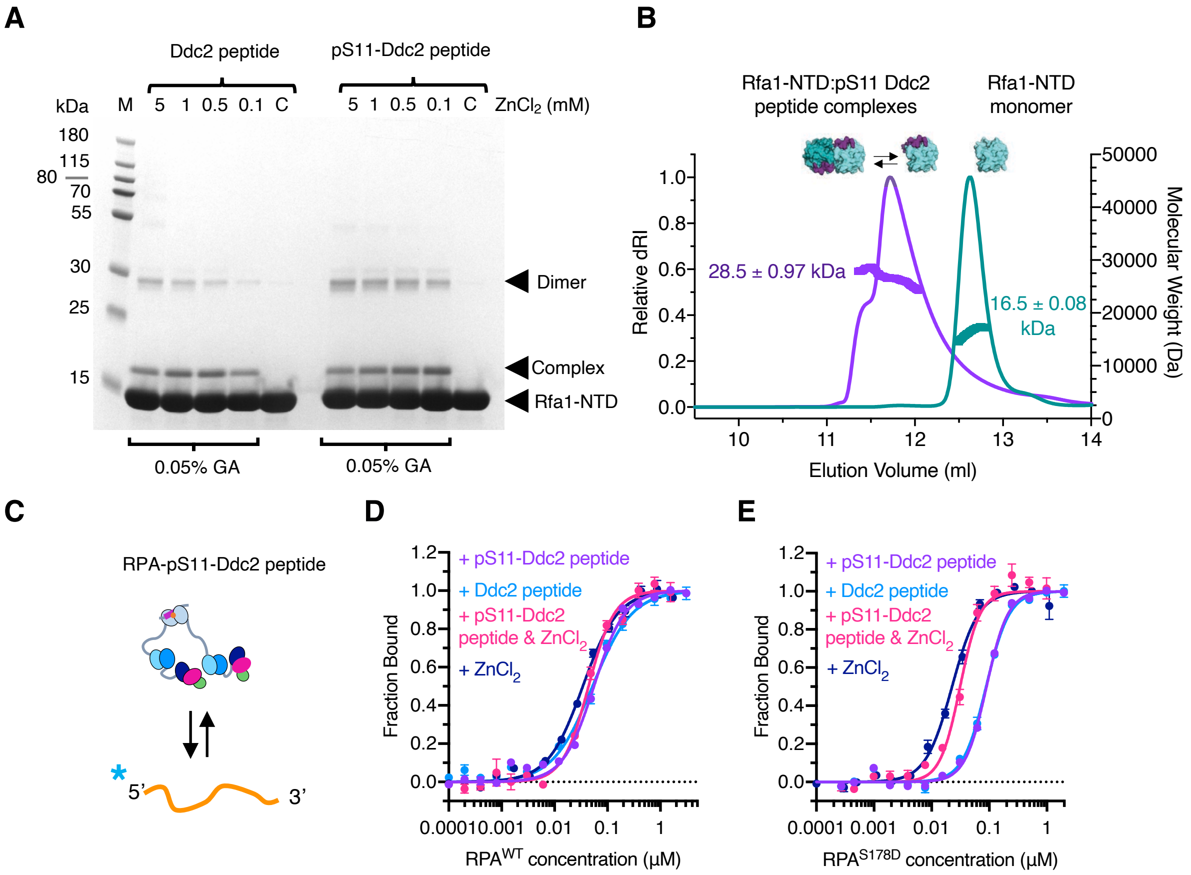


**F**

| RPA complex vs Cy5-dT$_{100}$ | EC$_{50}$ (nM) | Hill (*h*) |
|---|---|---|
| RPA$^{WT}$ + Ddc2 Peptide | 52 ± 5 | 1.2 ± 0.1 |
| RPA$^{WT}$ + pS11-Ddc2 Peptide | 56 ± 5 | 1.5 ± 0.2 |
| RPA$^{WT}$ + pS11-Ddc2 Peptide + ZnCl$_2$ | 44 ± 3 | 1.8 ± 0.2 |
| RPA$^{WT}$ + ZnCl$_2$ | 34 ± 2 | 1.3 ± 0.1 |
| RPA$^{S178D}$ + Ddc2 Peptide | 88 ± 3 | 2.1 ± 0.2 |
| RPA$^{S178D}$ + pS11-Ddc2 Peptide | 90 ± 3 | 2.3 ± 0.2 |
| RPA$^{S178D}$ + pS11-Ddc2 Peptide + ZnCl$_2$ | 32 ± 2 | 2.5 ± 0.3 |
| RPA$^{S178D}$ + ZnCl$_2$ | 23 ± 1 | 1.3 ± 0.2 |

**Supplementary Figure 8**

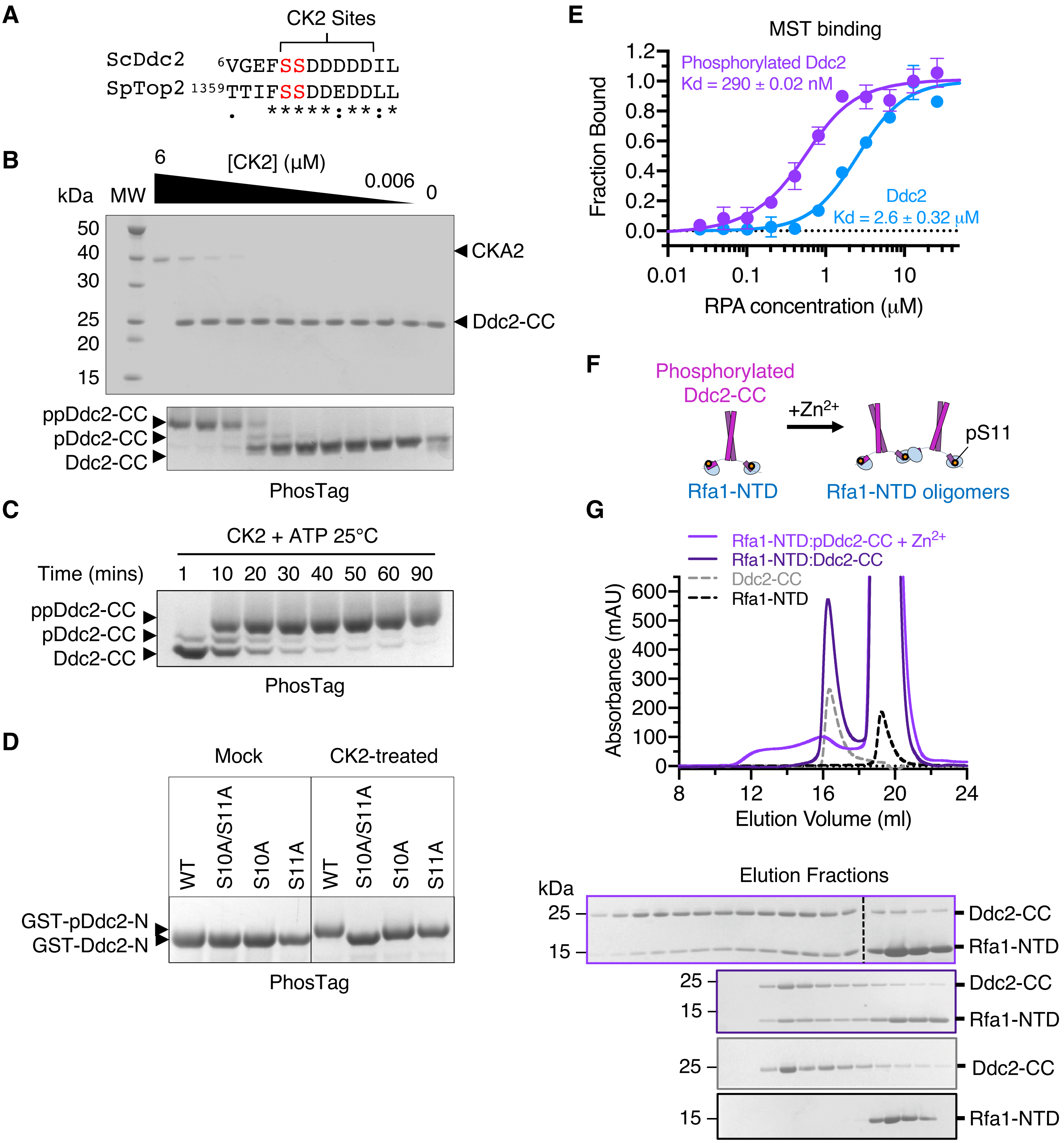


**Supplementary Figure 9**

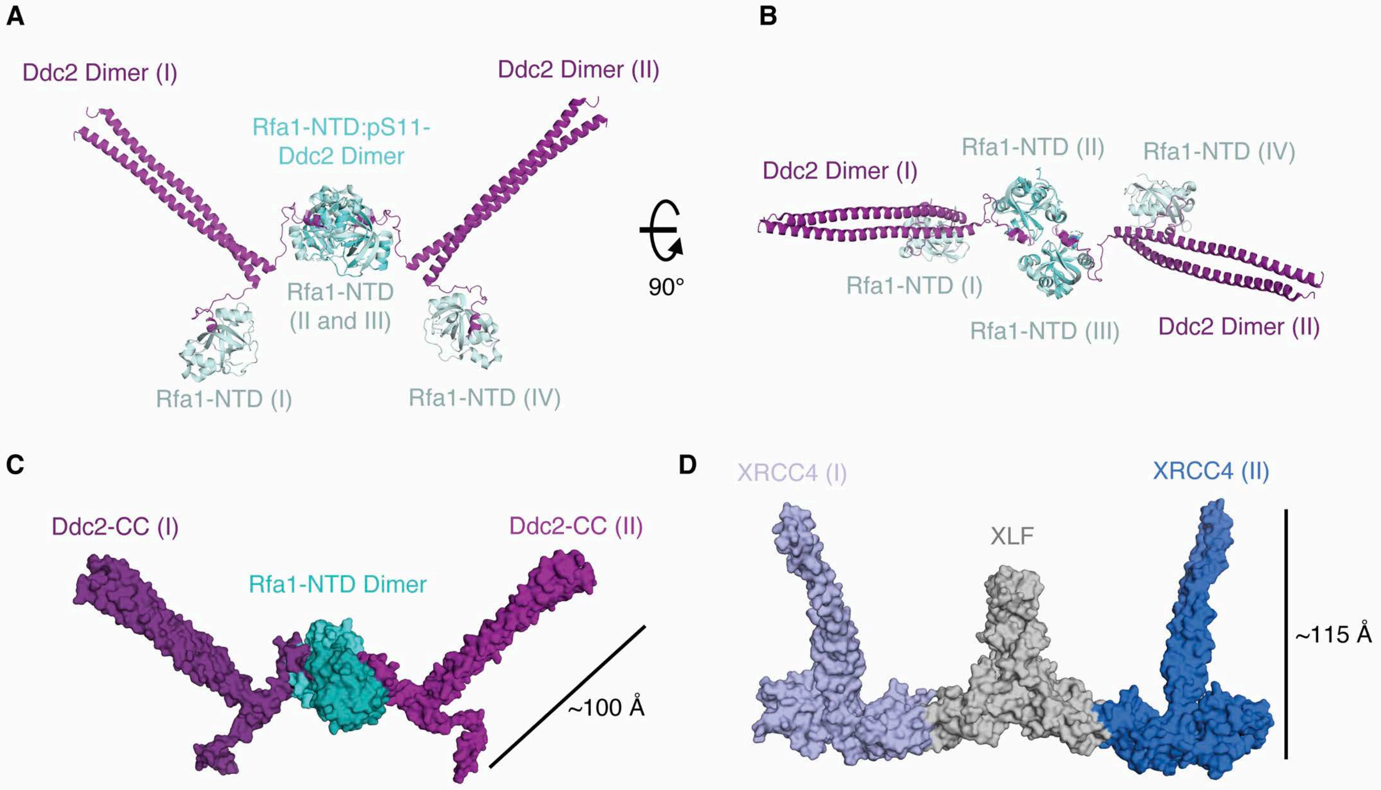


**Supplementary Figure 10**

### Supplementary Figure Legends

**Supplementary Figure 1 – Interaction studies between Mec1-Ddc2 and RPA.**
**A** Schematic of a Mec1-Ddc2 pull down using immobilised ssDNA ($dT_{32}$) coated with $RPA^{WT}$ or $RPA^{S178D}$.
**B** Corresponding SDS-PAGE analysis of the resultant protein samples after pull-down.

**Supplementary Figure 2 – Interaction between Ddc2-CC homodimer and RPA-ssDNA.**
**A** SDS-PAGE of purified recombinant Ddc2-CC.
**B** SEC-MALs of Ddc2-CC indicating that it is a dimer in solution (~20kDa monomer).
**C** MST binding of fluorescently labelled Ddc2-CC vs RPA complex.
**D** Electrophoresis mobility shift assay (EMSA) of RPA with 5'Cy5-$dT_{100}$ at different RPA:ssDNA ratios (assuming a 30nt binding site for RPA), with and without the addition of Ddc2-CC. The dimeric Ddc2-CC fragment stabilises RPA-$dT_{100}$ complexes, which suggests Ddc2-CC induces cooperative ssDNA binding of RPA. This is more evident when the ssDNA is undersaturated.
**E** Schematic of the RPA-ssDNA capture assay.
**F** Resulting Coomassie stained SDS-PAGE (upper gel) and in-gel Cy5 fluorescence (lower gel) to detect the Rfa1 and ssDNA, respectively. The fold difference is shown below and is the same for both protein and DNA using lane 2 as a reference.

**Supplementary Figure 3 – Mec1-Ddc2 and RPA protein samples for full-length interaction studies.**
**A** PhosTag SDS-PAGE analysis of Mec1-Ddc2 showing a mixed population of phosphorylated polypeptides. Ddc2 can be readily dephosphorylated by the addition of Lambda-PP.
**B-C** SDS-PAGE of protein samples used for interaction studies by MST. Mec1-Ddc2 was purified in the presence of phosphatase inhibitor cocktail (B-glycerophosphate, sodium fluoride, sodium pyrophosphate) to preserve phosphorylation. The best fractions from a Heparin step, based on the SDS-PAGE (boxed), were pooled and aliquoted into two volumes, one of which was treated with lambda-phosphatase and $MnCl_2$.

**Supplementary Figure 4**
**A** Growth of *DDC2* mutants. Strains PY434, PY436, and PY437 are *ddc2Δ* and contain plasmid p(*LEU2 ddc2-x*). *TEL1* and *DDC1* status is shown on the right. The strains were tested for growth on YPDA plates with or without hydroxyurea as indicated.
**B** Phosphorylation of Rad53 by Mec1-Ddc2 or Mec1-Ddc2x mutants with the indicated concentrations of Dpb11 activator. To eliminate contributions by Rad53 kinase itself, the kinase-dead version was used (Rad53-kd, K227A).
**C-D** Raw Isothermal Titration Calorimetry (ITC) and normalised heat of binding isotherms for Ddc2 pS10-pS11 phosphopeptides titrated against Rfa1-NTD (purple) or buffer alone (heat of dilution controls, grey).

**Supplementary Figure 5 – Rfa1-NTD purification and structure comparisons.**
**A-B** Final size exclusion chromatogram of Rfa1–NTD along with SDS-PAGE analysis of fractions from the major elution peak (inside the green box). Estimated molecular weight elution volumes for standard proteins are indicated (arrows) and labelled.
**C** Comparison between apo and pS11-Ddc2 peptide bound Rfa1-NTD solved in this study. Interacting amino acid side chains are shown for both apo and phosphopeptide complex.
**D** Structural comparison between Rfa1-NTD in complex with *S. cerevisiae* pS11-Ddc2 peptide (purple) and Rfa1-NTD in complex with the equivalent region of *K. lactis* Ddc2 (pink, PDB 5OMB). Structures were aligned on Rfa1-NTD.

**E** Structural superposition of Rfa1-NTD, or the equivalent human RPA70-NTD, in complex with peptides (p53, PDB 2B3G; PrimPol, PDB 58NA and 5N85; DNA2, 5EAY; ATRIP, PDB 4N3B). The structures were aligned on the OB-fold and the Saccharomyces cerevisiae Rfa1-NTD is shown for clarity.

**Supplementary Figure 6 – Ddc2 N-terminal peptide binds $ZnCl_2$ but not $CaCl_2$ or $MgCl_2$.**
**A** Coordinated metal ions in the Rfa1-NTD:pS11-Ddc2 peptide complex structure. Positive difference density is shown as green mesh.
**B-H** Raw Isothermal Titration Calorimetry (ITC) isotherms of metal solutions against Ddc2 peptide variants.

**Supplementary Figure 7 – MST of Ddc2 pS11 peptide interaction with $ZnCl_2$.**
**A-B** 6-FAM labelled Ddc2 pS11 phosphopeptide in complex with Rfa1-NTD binds $ZnCl_2$ with higher affinity than peptide alone. Consistent with ITC, calcium, or magnesium, does not bind, and other physiological metals (manganese, and iron) do not interact with the complex. MST curves were fitted with the law of mass action equation in Nanotemper analysis software and are listed in **b,** n.d. indicated not determined. A peptide only control binds $ZnCl_2$ with a very similar affinity (132 µM) to that obtained from ITC experiments (110 µM).

**Supplementary Figure 8 – Rfa1-NTD in complex with Ddc2 peptides dimerise in the presence of zinc.**
**A** SDS-PAGE analysis of Rfa1-NTD:Ddc2 peptide or Rfa1-NTD:pS11-Ddc2 peptide crosslinking (0.05% glutaraldehyde, GA) under differing $ZnCl_2$ concentrations (labelled above the lanes). A control, no crosslinker, sample was also included.
**B** Size exclusion chromatography (SEC) with inline multi-angle light scattering (MALS) of Rfa1-NTD:pS11-Ddc2 (purple) in the presence of 5mM $ZnCl_2$. The mass calculations suggest an equilibrium of species that approaches the molecular weight of a dimer and is clearly larger than the monomeric Rfa1-NTD alone (teal, predicted monomer MW 15.2 kDa).
**C** Schematic of Ddc2 phosphopeptides tethering RPA, based on the structure, to influence its association with ssDNA.
**D-E** MST-binding curves of RPA complexes interacting with Cy5-labeled $dT_{100}$ ssDNA substrate, using $RPA^{WT}$ or $RPA^{S178D}$, in the absence or presence of Ddc2 peptides/phosphopeptides and/or $ZnCl_2$ (1mM final). Normalized fluorescence was calculated using NanoTemper Analysis 1.2.101. Measurements were repeated at least 3 times or more and data points are averages with standard error (SE).
**F** For all MST curves, apparent $EC_{50}$ and Hill values were calculated using the Hill-equation and tabulated.

**Supplementary Figure 9 – CK2 phosphorylates Ddc2-CC in vitro.**
**A** sequence alignment between *S. cerevisiae* Ddc2 N-terminal region, containing phosphosites S10 and S11, and a known CK2 bisphosphorylation site in *S. pombe* Topoisomerase II, which shares similar motif.
**B** Phosphorylation assay of Ddc2-CC using a serial dilution of recombinant CK2 (ScCKA2) indicated. The reactions were resolved on SDS-PAGE (upper) and a PhosTag gel (lower).
**C** Time resolved CK2 phosphorylation assay resolved on PhosTag SDS-PAGE.
**D** PhosTag SDS-PAGE of purified Ddc2 N-terminal fragment (amino acids 1-37) fused to GST, with no (WT), one (S10, S11), or both (S10/S11) phosphosite substituted for alanine.
**E** MST-binding curves of labelled Ddc2-CC, treated with CK2 and repurified (phosphorylated), and Ddc2-CC alone against a titration of RPA complex.
**F** Schematic of Ddc2-CC:Rfa1-NTD complexes forming higher order oligomers in the presence of zinc upon phosphorylation.

**G** Size exclusion chromatography analysis of phosphorylated Ddc2-CC in complex with Rfa1-NTD. When $ZnCl_2$ is included (0.1mM) the complex spreads over an earlier elution volume range indicating a range of higher order oligomers.

**Supplementary Figure 10 – Models of higher order assemblies of Rfa1-NTD and Ddc2-CC dimers.**

**A-B** structural superposition of *S. cerevisiae* Rfa1-NTD in complex with *K. lactis* Ddc2 (PDB 5OMB) onto a dimer of Rfa1-NTD:pS11-Ddc2 determined in this study. This array does not produce clashes that would prohibit this assembly.

**C-D** Comparison of the Rfa1-NTD:Ddc2-CC higher order assemblies with XRCC4-XLF structures (PDB 7LT3).
